# Digenic genotypes: the interface of inbreeding, linkage, and linkage disequilibrium

**DOI:** 10.1101/2022.01.07.475385

**Authors:** Reginald D. Smith

## Abstract

Many traits in populations are well understood as being Mendelian effects at single loci or additive polygenic effects across numerous loci. However, there are important phenomena and traits that are intermediate between these two extremes and are known as oligogenic traits. Here we investigate digenic, or two-locus, traits and how their frequencies in populations are affected by non-random mating, specifically inbreeding, linkage disequilibrium, and selection. These effects are examined both separately and in combination to demonstrate how many digenic traits, especially double homozygous ones, can show significant, sometimes unexpected, changes in population frequency with inbreeding, linkage, and linkage disequilibrium. The effects of selection on deleterious digenic traits are also detailed. These results are applied to both digenic traits of medical significance as well as measuring inbreeding in natural populations.

## 1. Introduction

Starting in the early 1960s, population geneticists began building on the advances made understanding the dynamics of single loci subject to evolutionary forces to analyze the dynamics of multilocus systems with two or more loci. Early work on multilocus genetics included (Hogben, 1932) and (Li, 1953) on double homozygous recessive traits and (Haldane, 1949) in the first steps of understanding the effects of inbreeding at two loci. Digenic models were also essential in theories by Fisher and others on the evolution of dominance (Fisher, 1928a,b, 1929).

The dynamics of two loci subject to selection was a subject of early work by Kimura (Kimura, 1956) but was first developed into a general theory of the effect of selection on digenic traits by Lewontin & Kojima (Lewontin & Kojima, 1960). Their paper described the general equations for the evolution of haplotype frequencies and the effects of linkage and linkage disequilibrium on changes in haplotype frequency and stable states. Key to the work by both groups of authors was the analysis of the dynamics of two loci where the relative fitness matrix is symmetric allowing the problem to be analytically tractable and guaranteeing the existence of stable solutions under certain conditions of relative fitness.

Lewontin and Kojima’s paper stimulated much interest in the topic, especially in light of the increasing availability of polymorphism data, which include (Bodmer & Felsenstein, 1967; Karlin & Feldman, 1970; Karlin, 1975, 1979; Karlin & Avni, 1981; Bürger, 2020). A good review of some of the historically important results can be found in (Karlin, 1975) and (Bürger, 2020).

### 1.1. Medical genetics and digenic traits

One of the first practical applications of the population genetics of two loci was in medical genetics. From the late 1970s to the 1990s, digenic traits, especially where they could possibly be of relevance to explaining medical conditions, were increasingly investigated. A disease model based of the symmetric fitness models of Lewontin, Kojima, and Karlin was given by (Merry et. al., 1979). In addition, the enumeration of medically relevant two locus genotypes and the covariance of relatives where the loci are unlinked is given by (Neuman & Chakravarti, 1992) building on earlier work by (Hartl, 1968). Full enumeration of all possible digenic trait genotypes were given by (Li & Reich, 2000; Hallgrímsdóttir & Yuster, 2008).

Two locus disease models continued to evolve throughout the 1980s and 1990s with important contributions by (Hodge, 1981a; Hodge & Spence, 1981b; Goldin & Weeks, 1993). These were complimented by the first confirmed discovery of a digenic disease trait: a double heterozygous genotype that causes a variation of retinitis pigmentosa (Kajiwara et. al., 1994). However, much interest in digenic traits temporarily waned with the advent of advanced technologies such as next generation sequencing and methods such as GWAS which allowed analysis of large numbers of loci compared to previously limited polymorphism data.

However, there has been a resurgence of interest in so-called oligogenic disorders, defined as disorders whose aetiology is described by the genotypes at two or more loci but whose primary mechanisms are epistatic and not additive. Oligogenic disorders occupy a midpoint between single locus Mendelian disorders and complex diseases defined as quantitative traits. Several reviews (Badano & Katsanis, 2002; Cooper et. al., 2013; Schäffer, 2013; Deltas, 2018) have discussed the increasing number of discoveries of digenic or oligogenic diseases. In (Schäffer, 2013) ninety-five reports of oligogenic disorders were listed along with the method of inheritance for each locus. In the Online Mendelian Inheritance in Man (OMIM) database (OMIM, 2022), there are currently 33 entries for disorders that have an either digenic dominant or digenic recessive aetiology. About half are inherited in an autosomal dominant manner with double heterozygote genotypes most commonly described. This however, is still minuscule compared to the over 2,800 disorders with autosomal recessive aetiology and over 2,000 disorders with autosomal dominant aetiology in OMIM. While still relatively small in confirmed numbers, they are subject of renewed search and interest especially in investigations of disorders with incomplete penetrance across a known disease genotype.

### 1.2. Measuring inbreeding using digenic genotypes

One of the most well-known effects of inbreeding is the increase of homozygosity and the decrease of heterozygosity at individual loci. However, when two locus dynamics are considered, some additional and unexpected effects emerge. As will be described in detail later, one effect of inbreeding is to cause correlations between genotypes at different loci, particularly when they are linked. This correlation is positive for two loci with homozygous genotypes or two loci with heterozygous genotypes. This has been exploited in fields such as genetics of wild populations or conservation genetics (David, 1998; Balloux et. al., 2004; David et. al., 2007) as a way to measure the extent of inbreeding in populations using genetic data where pedigrees are unavailable. This provides an alternate measure of inbreeding besides other measures such as runs of homozygosity.

### 1.3. Applications of the dynamics of digenic traits

While much rich theoretical work has been done on two locus systems over the past 50 years, the bulk of work has been restricted to the theoretical realm, much more so than either single locus or polygenic population genetics. One of the most interesting aspects has been the lack of attention to the effects of selection on favorable or deleterious digenic traits which are driven to fixation or extinction. The majority of theoretical studies on selection and digenic traits have been the studies of stable equilibria. Similarly, while the effects of linkage disequilibrium and non-random mating are known, they have only occasionally entered the applied genetics realm, possibly due to a relative lack of expositions that clearly apply theoretical results to applications of interest in medical or conservation genetics. The purpose of this paper is to not only review past results but allow them to be put into a context where they are more accessible to the applied genetics field and can be used to understand or interpret results in the burgeoning study of oligogenic traits, particularly deleterious ones. This will enable a greater appreciation for the effects of inbreeding, linkage disequilibrium, and selection for digenic traits which is common for Mendelian or complex traits.

## 2. The effects of inbreeding on two loci in linkage equilibrium

It has been known for centuries that breeding of relatives can lead to an increasing frequency of relatively rare disorders. The modern analysis of inbreeding was formalized by Wright (Wright, 1922) and improved by Malécot’s probabilistic definition of the probability of two alleles at a locus being identical by descent (IBD) (Malécot, 1970). The inbreeding coefficient of descent *F* is defined as the IBD probability for two alleles in a locus and is equal to the coefficient of coancestry, *θ*, of the parents, also called kinship (e.g. (Jacquard, 1975)). In other words, the inbreeding of the progeny reflects the consanguineous relationship of the parents. Therefore the probability, *F*, that two alleles in the progeny are IBD is equal to the probability that two alleles from the same locus in each parent are IBD as well.

The genotypic results of inbreeding are widely known, most markedly increased homozygosity by an amount *p*(1 – *p*)*F* and a decline in heterozygosity. This increase in homozygosity allows the appearance of rare homozygous genotypes at significantly higher frequencies than if the population was outbred. This increases the frequencies of recessively inherited disorders in inbred populations.

The story for two loci is a bit more complex. First investigated by JBS Haldane (Haldane, 1949) and most thoroughly expounded in the collaboration of C. Clark Cockerham and Bruce Weir (Weir & Cockerham, 1968;

Cockerham & Weir, 1968, 1973, 1977; Weir & Cockerham, 1974), the effects of inbreeding across two loci involves not just the effects of increased homozygosity on the joint occurrence of genotypes.

Where two loci are concerned, we want to first investigate the conditions under which alleles at each locus in different individuals are identical by descent. We will designate the two alleles at the first locus in individuals 1 and 2 as *a*_1_ and *a*_2_ and at the second locus as *b*_1_ and *b*_2_. Note that these only represent the *identity* of the alleles and not the specific allele present. Per the notations of Weir and Cockerham we will define four different types of two-locus inbreeding coefficients:

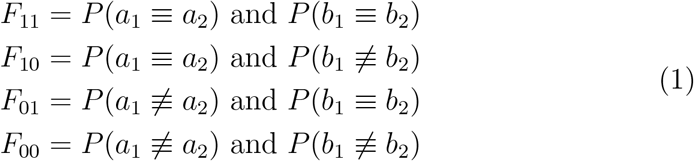

The ≡ symbol indicates identity by descent. Note the single locus inbreeding coefficient for locus *a* is *F*_1_. = *F*_11_ + *F*_10_ and the coefficient for locus *b* is *F*_.1_ = *F*_11_ + *F*_01_. The average of these two is often used to represent the average probability of IBD across both loci and is designated as *F*_1_ = (*F*_1_. + *F*_.1_)/2. For our analysis, we will assume the IBD probabilities at each locus are identical so *F*_.1_ = *F*_1._ = *F*_1_ = *F*. In evaluating the effects of inbreeding on two loci, the coefficient *F*_11_ is most useful in describing the impact of inbreeding at both loci. A graphical representation of *F*_11_ is given in Figure 3.

**Figure 1:**
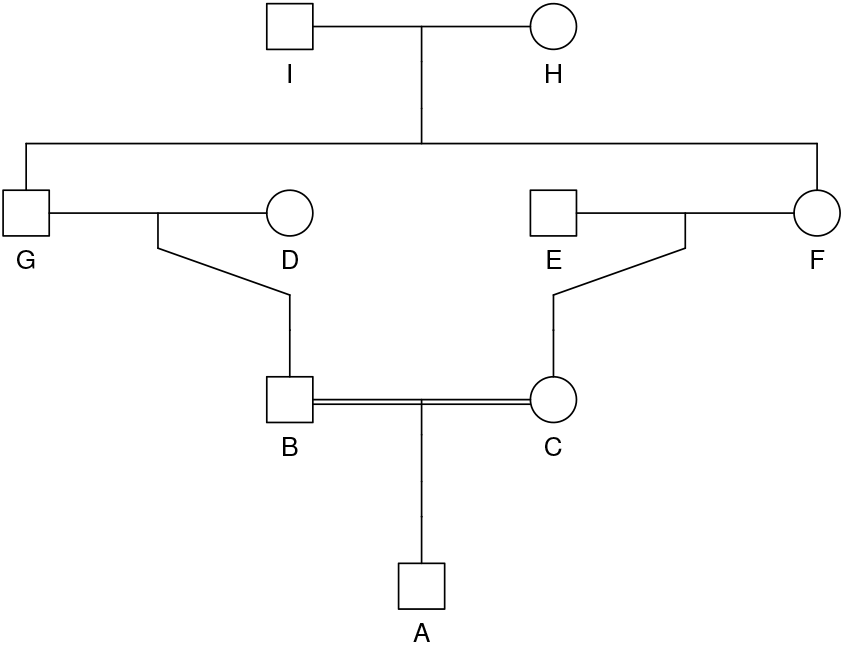
Pedigree of first cousin inbreeding.

**Figure 2:**
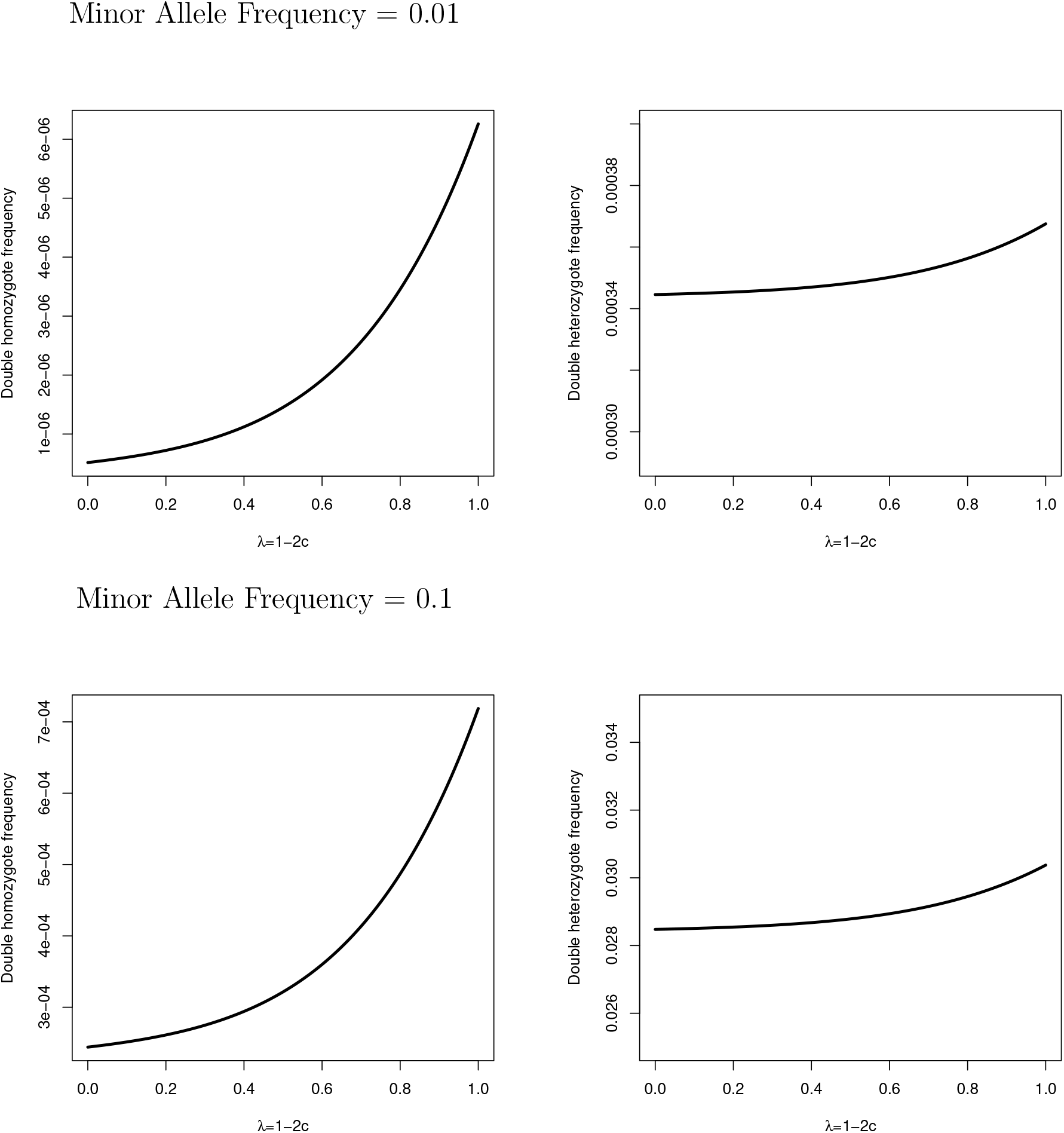
Plots of the frequencies of double homozygous and double heterozygous genotypes where the minor allele *q* at both loci is 0.01 (top graphs) and 0.1 (bottom graphs) versus linkage designated by the Schnell linkage value λ = 1 – 2*c*.

**Figure 3:**
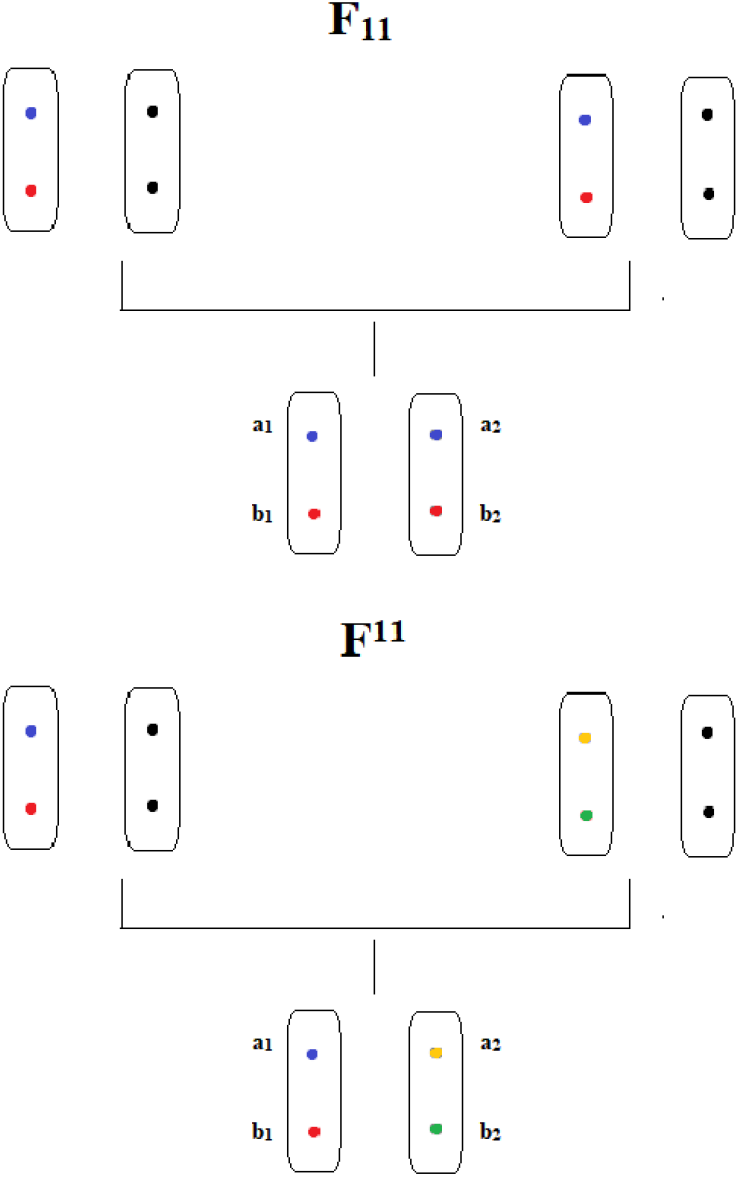
Images representing *F*_11_ and *F*^11^ based on ancestral chromosome haplotypes and the haplotypes of descendants. All identically colored non-black colored alleles are identical by descent. For all diagrams, the two pairs of chromosomes are arbitrary paternal and maternal ancestors (which can be one individual sharing both chromosomes) and the bottom chromosome pair is a single descendant.

In particular for the determination of genotype frequencies is the importance of the identity disequilibrium (ID), *η*_11_. The ID is a measurement of the increased frequency of joint IBD alleles at both loci above the IBD expected based on the product of the inbreeding coefficients at each locus. Where the inbreeding coefficient is the same for both loci, ID is defined as

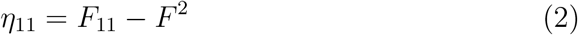

The coefficient *F*_11_, however is not a fixed value across all loci pairs and all individuals in the population. In particular, *F*_11_ depends on the level of linkage between the two loci as well as the pedigree of ancestors back to the original gamete to account for all possibilities of transmission. Its boundary values are *F*_11_ = *F*^2^ if *c* = 1/2, where *c* is the recombination frequency, and there is no linkage. Thus for *c* = 1/2, *η*_11_ = 0. For complete linkage where *c* = 0, *F*_11_ = *F* and thus *η*_11_ = *F*(1 – *F*). Intermediate values of *F*_11_ are calculated with algorithms given the pedigree of the individual. These will be explained in Appendix A but are introduced in (Haldane, 1949; Weir & Cockerham, 1968).

The main effect of identity disequilibrium generated by inbreeding is to increase the frequency of double homozygous and double heterozygous genotypes while reducing the frequency of genotypes that are pairs of homozygous and heterozygous genotypes. The increase in double homozygotes under inbreeding is largely expected but a positive counterbalance to the decrease of double heterozygous genotypes under inbreeding is often not appreciated. The effects of *η*_11_ by genotype are shown in Table 1. One way to interpret identity disequilibrium is as the correlation between genotypes at different loci. In other words, two loci are more likely to show double homozygosity or double heterozygosity than expected by chance.

**Table 1:** Two loci genotype frequencies under inbreeding at linkage equilibrium taking identity disequilibrium into account. For original explanations and derivations see (Haldane, 1949), though he uses *ϕ* for *η*_11_. The variance of each locus is represented by 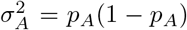 and 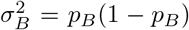. The variable *η*_11_ is the identity disequilibrium. Per the regular results of inbreeding, 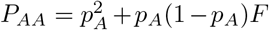, *P_Aa_* = 2*p*_A_(1 – *p_A_*)(1 – *F*), *P_aa_* = (1 – *p_A_*)^2^ + *p_A_*(1 – *p_A_*)*F*, 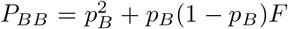, *P_Bb_* = 2*p_B_*(1 – *p_B_*)(1 – *F*), *P_bb_* = (1 – *p_B_*)^2^ + *p_B_*(1 – *p_B_*)*F*. If the loci are unlinked *η*_11_ = 0 and if they are completely linked *η*_11_ = *F*(1 – *F*). Note double heterozygotes have a factor of four for identity disequilibrium due to the two types of combining gametes (*AB* and *ab* or *Ab* and *aB*) that create double heterozygotes and the possibility of each gamete being on the paternal or maternal contributed chromosome. Similarly homozygotes-heterozygotes have a factor of two in relation to identity disequilibrium since each genotype can appear two different ways depending on its contribution from maternal or paternal chromosomes.

| | $AA$ | $Aa$ | $aa$ |
| --- | --- | --- | --- |
| $BB$ | $P_{AA}P_{BB} + \sigma_A^2\sigma_B^2\eta_{11}$ | $P_{Aa}P_{BB} - 2\sigma_A^2\sigma_B^2\eta_{11}$ | $P_{aa}P_{BB} + \sigma_A^2\sigma_B^2\eta_{11}$ |
| $Bb$ | $P_{AA}P_{Bb} - 2\sigma_A^2\sigma_B^2\eta_{11}$ | $P_{Aa}P_{Bb} + 4\sigma_A^2\sigma_B^2\eta_{11}$ | $P_{aa}P_{Bb} - 2\sigma_A^2\sigma_B^2\eta_{11}$ |
| $bb$ | $P_{AA}P_{bb} + \sigma_A^2\sigma_B^2\eta_{11}$ | $P_{Aa}P_{bb} - 2\sigma_A^2\sigma_B^2\eta_{11}$ | $P_{aa}P_{bb} + \sigma_A^2\sigma_B^2\eta_{11}$ |

Where inbreeding is present, the expected single-locus homozygous and heterozygote frequencies are given in the text of Table 1. Measuring covariance similar to that in linkage disequilibrium, we can show the covariances for double heterozygosity and double homozygosity at loci are only based on the identity disequilibria and the variances of heterozygosity frequency at each locus.

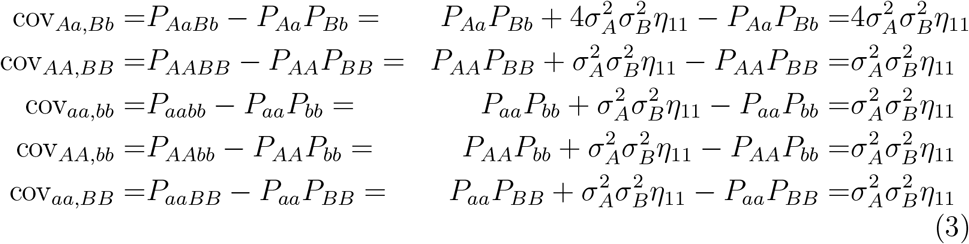

The correlations, however, are different though positive in these cases as shown below. A key finding is that the strength of the correlations varies not only with the identity disequilibrium but also directly with the degree of polymorphism of the loci. The larger the variance of each locus, the larger the correlation. As *F* increases towards 1, however, *F*_11_ and *F* both approach one (see (Cockerham & Weir, 1973), page 316) since the widespread homozygosity reduces the effect of recombination to create different varieties of genotypes in offspring and so the correlation approaches zero equaling zero when *F* = 1.

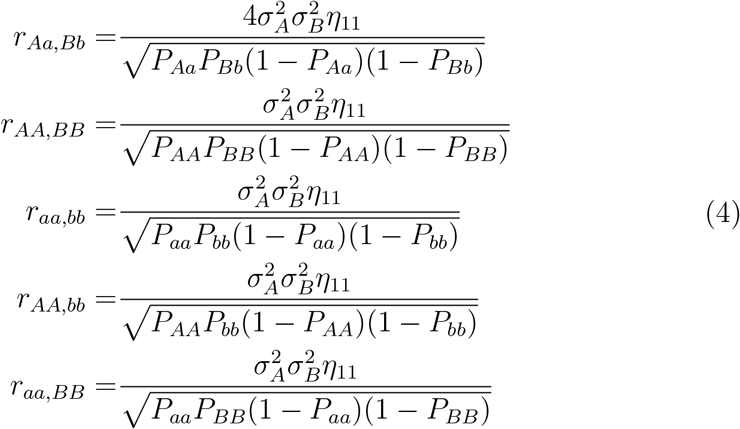

## 2.1. Using identity disequilibrium to estimate selfing and measure inbreeding

One of the first practical uses for identity disequilibrium was given by (David et. al., 2007). Using previous results from (Cockerham & Weir, 1973b) that showed that identity disequilibrium could exist at unlinked loci when the organism was capable of self-fertilization, the authors of (David et. al., 2007) devised a metric *g*_2_ that used the double heterozygosity frequency across many pairs of loci to approximate *η*_11_. Of importance was the fact that this was applicable only to those organisms who could self-fertilize: mostly plants but also some animals such as the freshwater snail *Bulinus truncatus*. Double heterozygosity frequencies could then be compared to heterozygosity frequencies at individual loci to estimate s, the percent of the population that was selfing. While *g*_2_ is both useful and well-characterized statistically, subsequent studies often departed from the selfing requirement and estimated *g*_2_ to determine if identity disequilibrium, and thus inbreeding, existed in populations where no selfing occurred. A meta-analysis of many of these studies (Miller & Coltman, 2014) found that *g*_2_ estimates were often not significantly different from zero. This is an issue not due to the number of markers, as some suspected, but rather the assumptions that unlinked loci can easily estimate identity disequilibrium in populations with no selfing.

As stated earlier, for unlinked loci *η*_11_ = 0 for pedigree mating in non-selfing populations. It is possible though for average values of *F*_11_ – *F*^2^ to exceed zero for unlinked loci where there is random mating in finite inbred populations. This was demonstrated in (Cockerham & Weir, 1969) (see discussion) where they showed that in finite populations where the population averages of the descent coefficient and inbreeding coefficient are given by 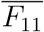 and 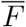

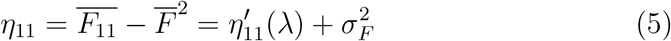

The variable 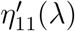 is the former definition of identity disequilibrium based on pedigree mating and linkage. The dependent parameter λ is the Schnell measure of linkage λ = 1 – 2*c* (Schnell, 1961) that ranges from [0, 1] as c ranges from [1/2, 0]. The variance 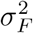 represents the variance in the measure of inbreeding across the same locus in the finite population that exists due to finite sampling effects. When the two loci are unlinked, λ = 0 and 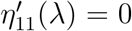 but the component 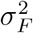 remains. However, in (Cockerham & Weir, 1969) it is shown through numerical calculation and plotting that 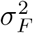 is normally of a near imperceptible magnitude and thus, *η*_11_ is still not likely to be much different from zero when the loci are unlinked.

Therefore it is important to state that *g*_2_, calculated amongst mostly unlinked loci, is not useful in determining inbreeding in populations where no selfing occurs. In (Cockerham & Weir, 1973b) (first equation on page 259), it is indicated that for unlinked loci, *η*_11_ = 0 when *s* = 0. There are, however, alternate ways that double heterozygosity can be used to infer the presence of inbreeding.

### 2.1.1. g_2_ vs. linkage regression

Given that *η*_11_ ≈ 0 for unlinked loci and *η*_11_ = *F*(1 – *F*) for complete linkage, the measurement of inbreeding using identity disequilibrium measures sure as *g*_2_ must account for linkage in non-selfing populations. This requires the use of loci that have linkage and will likely require a more dense set of markers in the genomes under analysis, with many markers occupying the same chromosomes required to analyze the changes with linkage. There should be a positive dependence of *g*_2_ on linkage, measured for example by regression, if inbreeding is present. This is complicated though by the effects of linkage disequilibrium that can be larger at linked loci. Correcting this will be addressed later in the paper.

## 2.2. Double homozygosity and double heterozygosity from first cousin matings

As discussed earlier, rare genetic disorders will cluster more in families or populations where consanguineous matings are more common. Consanguineous matings can be a significant portion of the population, even in countries where culture and legal environments mitigate against it. For example, using UK Biobank data (Yengo et. al., 2019) showed that about 1 out of every 3,652 individuals showed extreme inbreeding as measured by runs of homozygosity. Here we will show the effect of first cousin matings on the expected occurrence of double homozygous and double heterozygous genotypes. This is an important angle where the medical genetics of rare disorders is concerned.

First derived in (Haldane, 1949), but also derived in Appendix A, the two locus descent coefficient *F*_11_ for the progeny of first cousins is

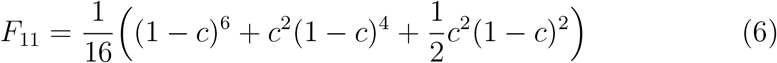

Given that *F* = 1/16 for first cousins as well, the identity disequilibrium is given by

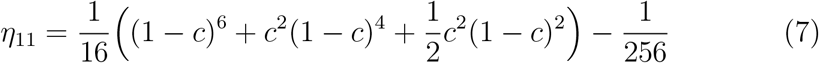

It is clear when *c* = 1/2, *η*_11_ = 0 and likewise when *c* = 0, *η*_11_ = *F*(1 – *F*) = 15/256. Per Table 1 when the loci are unlinked the first cousin digenic genotypes are the products of the genotypes at each locus under inbreeding. With linkage, however, the frequencies of double homozygous and double heterozygous genotypes begin to increase. This is particularly important for double homozygous genotypes since both the single and dual locus genotypes increase under inbreeding. Double heterozygote genotypes increase with linkage, however, the effect of identity disequilibrium is to reduce the rate of double heterozygosity decline due to the decline of heterozygosity at single loci and thus is more limited.

In Figure 2, the increase in double homozygotes and double heterozygotes are shown for increasing linkage where the minor allele frequencies are 0.01 or 0.1 in each case. The x-axis uses λ in order to show increasing linkage using an increasingly positive variable.

It is clear that the double heterozygous genotypes show little increase with linkage while double homozygous genotypes increase by over an order of magnitude with increasing linkage when the minor allele as a frequency of 0.01. Without inbreeding, the double homozygous frequency would only be 1 × 10^−8^ or 1 in 100 million. For first cousins this increases to 5.2 × 10^−7^ (1 in 1.9 million) when the loci are unlinked. For full linkage, however,this increases to 6.3 × 10^−6^ about 0.63 cases per 100,000 first cousin births. Granted these probabilities are calculated across the population of progeny of full cousins only but are still a remarkable increase in likelihood for such rare disorders.

This has implications for the populations in which digenic genotypes, especially those of medical interest occur. Double heterozygous genotypes do not see a significant increase frequency with linkage and the heterozygosity at the individual loci is also reduced by inbreeding. Therefore, autosomal dominant transmitted disorders of digenic character are likely more prevalent in the general population than consanguineous families, regardless of linkage. In the review of then known oligogenic disorders (2+ loci) (Schäffer, 2013), 48 of the 95 then published digenic maladies were transmitted in an autosomal dominant manner at both loci. In almost all of the individual reports, the genotypes reported are double heterozygous and most occurred in unrelated families and in only 13 of the disorders where the loci linked.

In contrast, double homozygous genotypes see large increases in frequency due to linkage and consanguineous matings. Of the 8 disorders recognized as having an autosomal recessive transmission at both loci in (Schäffer, 2013), 3 of 8 had some linkage and 5 of 8 looked at cases where the parents were consanguineous. This emphasizes the numerical results above that for rare variants at two loci, double homozygous genotypes are extremely rare in the absence of inbreeding and made more frequent by linkage.

## 3. The effects of inbreeding on two loci in linkage disequilibrium

Having discussed the frequencies of digenic genotypes when both loci are in linkage equilibrium, we can now address the more complex situation when the loci are in linkage disequilibrium and the genotype distributions reflect the effects of inbreeding. While linkage disequilibrium is ubiquitous, certain demographic processes make it more widespread in populations. In particular, population admixture and population bottlenecks significantly increase the linkage disequilibrium amongst loci over large sections of the genome.

### 3.1. Demographic process that generate linkage disequilibrium

One of the key aspects of digenic traits separating them from monogenic ones is the importance of both linkage and linkage disequilibrium. The frequencies of digenic genotypes and the effects of mating systems on those frequencies are heavily dependent on these variables. Given this fact, populations that have large linkage disequilibrium should be expected to have impacts on the frequencies of digenic traits, especially when mating between relatives occurs.

It has been long established that many demographic processes can generate or reduce linkage disequilibrium. Population admixture and population bottlenecks can generate linkage disequilibrium amongst loci while rapid population growth can reduce it (Slatkin, 1994). Linkage disequilibrium creation will be briefly reviewed for these two processes in the next section. Here linkage disequilibrium *D* and it’s normalized version, *D*′ are defined according to the standard notations for alleles *A* and *B* and different loci.

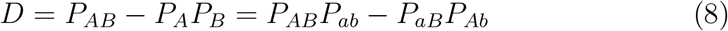

For *D*′ where *D* > 0

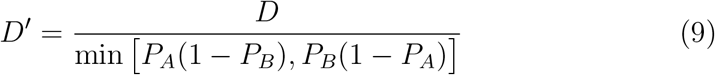

and where *D* < 0

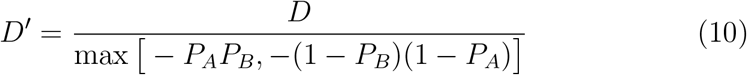

#### 3.1.1. Linkage disequilibrium from population admixture

Admixture between populations with different allele frequencies at a pair of loci will generate linkage disequilibrium between these loci, even if it is absent in the original populations. Though this linkage disequilibrium decays over time unless maintained by evolutionary forces, it can last for a substantial number of generations in cases where the loci in question are linked and encounter recombination with much lesser frequency.

The amount of linkage disequilibrium generated by admixture was first worked out by (Nei & Li, 1973). For the case of bi-allelic loci where population one contributes a proportion *m*_1_ to the final population this leads to

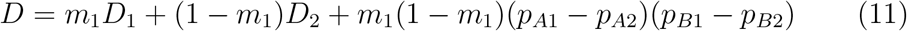

The *D*_1_ and *D*_2_ terms represent initial linkage disequilibrium in the original populations and the probabilities are the frequencies of allele *A* and *B* in population 1 or 2. This linkage disequilibrium affects haplotype frequencies which in turn help determine digenic trait frequencies.

#### 3.1.2. Linkage disequilibrium from population bottlenecks

A bottleneck is generally defined as a drastic reduction in effective population size. This reduction in population size allows drift to become a much stronger force leading to a reduction in genetic variation, increase in homozygosity, and linkage disequilibrium that is generated from the effects of drift and balanced by recombination (Hill & Robertson, 1968). As long as the population remains small, the linkage disequilibrium can be maintained since it is continuously generated by drift. This is in contrast to linkage disequilibrium by admixture which is a one-time even unless the admixture is continuous. However, as the population grows, the linkage disequilibrium will decline as the effects of drift weaken and recombination becomes dominant (Slatkin, 1994).

### 3.2. The effects of linkage disequilibrium on digenic genotype frequencies

Linkage disequilibrium has long been known to generate genotypes with an excess of homozygosity for two locus genotypes. Some measures of linkage disequilibrium explicitly take this homozygosity into account to measure linkage disequilibrium across groups of multiple loci (Sabatti & Risch, 2002; Rosenberg & Blum, 2007). In the next sections, we will outline the separate effects on digenic genotype frequencies due to linkage disequilibrium and inbreeding. First we will look at the basic effects of linkage disequilibrium on two locus genotype frequencies in outbred populations. Second we will introduce new descent coefficients required to understand the effect of linkage disequilibrium at two loci when inbreeding occurs followed by detailed calculations of genotype frequencies. Finally, we will revisit the first cousin mating example and discuss the measurement of linkage disequilibrium when Hardy-Weinberg equilibrium is not present.

#### 3.2.1. Digenic genotypes in outbred populations

When the population is outbred, the effect of linkage disequilibrium on genotype frequencies is clear as shown in Table 2. For positive linkage disequilibrium between loci, the frequencies of some genotypes can increase markedly. This alone can increase the frequency of digenic genotypes in recently admixed populations.

**Table 2:** Two loci genotype frequencies as a function of linkage disequilibrium absent inbreeding. See also (Lewontin & Kojima, 1960; Weir, 2008). Chart adopted from (Weir, 2008). *P_AB_* = *p_A_p_B_* + *D*, *P_ab_* = (1 – *p_A_*)(1 – *p_B_*) + *D*, *P_Ab_* = *p_A_*(1 – *P_B_*) – *D*, *P_aB_* = (1 – *p_A_*)*p_B_* – *D*. Note the double heterozygote (*AaBb*) genotype has two terms due to the genotype arising from either the dual gametes *AB* and *ab* or *Ab* and *aB*.

| | $AA$ | $Aa$ | $aa$ |
| --- | --- | --- | --- |
| $BB$ | $P_{AB}^2$ | $2P_{AB}P_{aB}$ | $P_{aB}^2$ |
| $Bb$ | $2P_{AB}P_{Ab}$ | $2P_{AB}P_{ab} + 2P_{Ab}P_{aB}$ | $2P_{aB}P_{ab}$ |
| $bb$ | $P_{Ab}^2$ | $2P_{Ab}P_{ab}$ | $P_{ab}^2$ |

**Table 3:**
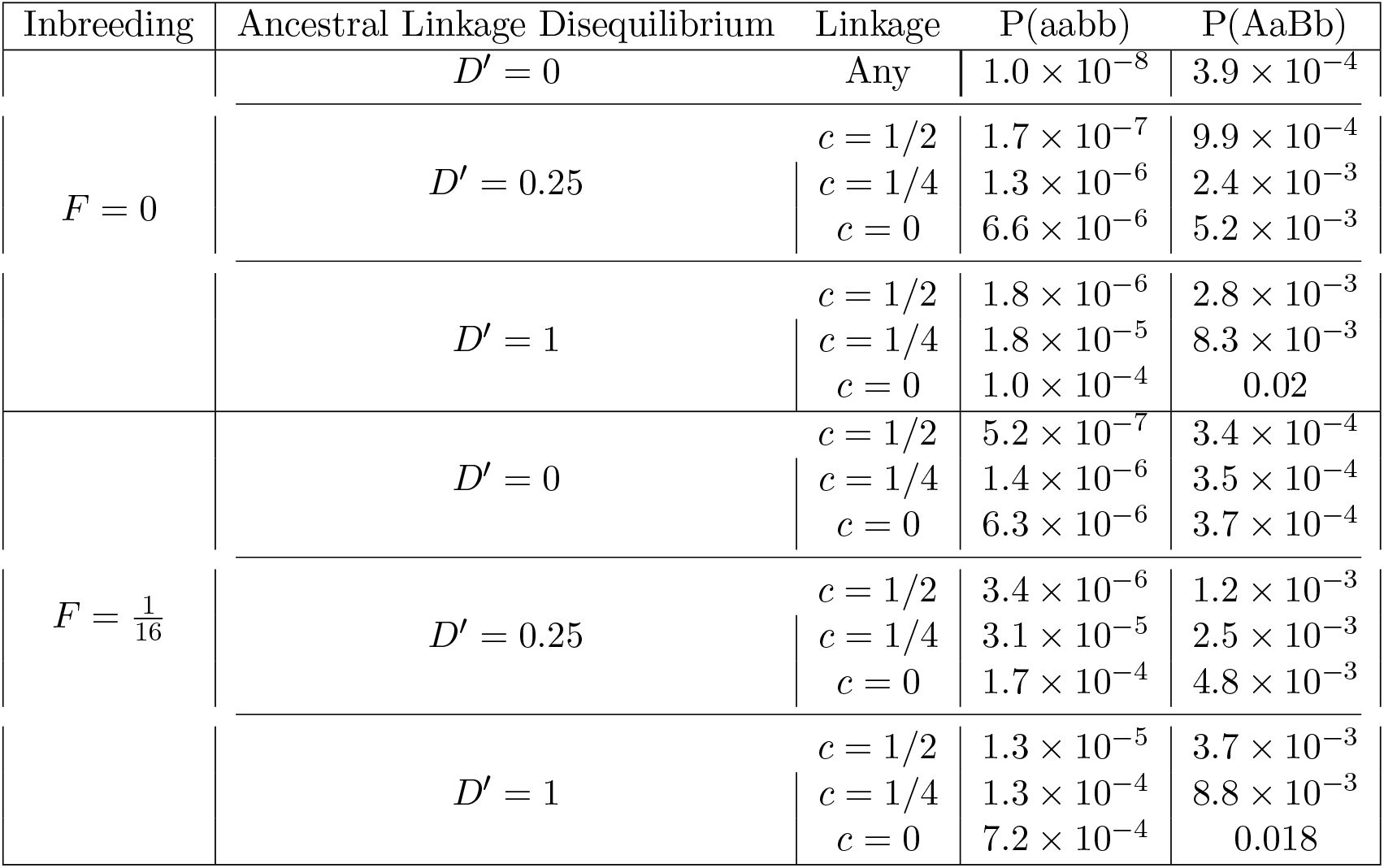
Expected double homozygote and double heterozygote frequencies under various combinations of ancestral linkage disequilibrium and inbreeding for *p_a_* = *p_b_* = 0.01 and consanguineous mating between first cousins determining *F* and the descent coefficients for the inbred case. Note that the final linkage disequilibrium in the affected population has undergone three generations of recombination from the ancestral linkage disequilibrium and in each case is *D*′(1 – *c*)^3^.

For example, for two loci with minor allele frequencies of 0.01, the expected double homozygous genotype frequency for the minor alleles is 1 × 10^−8^. With linkage disequilibrium equivalent to *D*′ = 0.25, however, this can increase to 6.6 × 10^−6^, over a 600-fold increase though still relatively uncommon.

#### 3.2.2. The combined impacts of inbreeding and linkage disequilibrium

The sole addition of identity disequilibrium to the genotype frequencies described in Table 1 assumes that the loci are in linkage equilibrium. This greatly simplifies the analysis. The presence of linkage disequilibrium changes frequencies due to the fact initial linkage disequilibrium interacts with linkage to change the expected frequencies of alleles being IBD within loci. It also means we have to account for several additional descent coefficients.

Linkage disequilibrium greatly increases the complexity of the analysis to an extent that cannot be covered fully in this paper. For detailed descriptions on the coefficients and their relationships in pedigree analysis, the author refers readers to (Cockerham & Weir, 1973, 1977). Their techniques for calculating coefficients with known pedigrees is also given in Appendix A. A key result though is linkage disequilibrium combined with inbreeding increases the frequency of double homozygous and double heterozygous genotypes in an additive fashion. In other words, there is an additive increase in the covariance between genotypes under inbreeding when linkage disequilibrium is present.

The first new coefficient, the parental descent coefficient *F*^11^, measures the probability that both of the gametes are identical to the gamete in the common ancestor both alleles descended from. In other words, *F*^11^ measures the probability that both gametes in each individual are passed down from the common ancestor without recombination.

Assuming ≡ means the alleles are from the same ancestral gamete, we can define the two locus parental coefficients similar to the inbreeding coefficients.

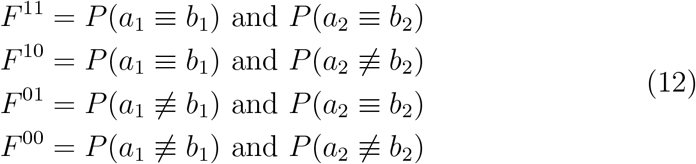

The average parental coefficient across both individuals can be defined similarly to the inbreeding coefficient case and is designated as *F*^1^. The coefficient *F*^11^ is graphically represented in Figure 3.

The second new coefficient, the recombination descent coefficient _11_*F* is a measure of the probability that an allele at the first locus on one gamete in one individual and an allele at the second locus on the other gamete in the same individual were originally part of the same gamete in the common ancestor. This measures the probability that there was a recombination event in one or both of the common parents and the alleles in the final generation are on different gametes than they started out. For more details see (Cockerham & Weir, 1973, 1977).

Similar to the definitions above for the parental coefficients

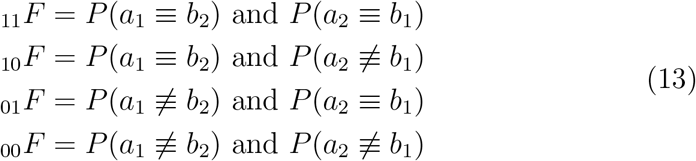

The ≡ symbol here indicates the two alleles at different loci on different gametes were originally together on the common ancestral gamete. These descent coefficients are necessary since descent identity no longer relies on alleles at a single locus but the shared gametic ancestry (or lack thereof) of the alleles on the gametes inherited by descendants. The coefficient 11_*F*_ is graphically represented in Figure 4.

**Figure 4:**
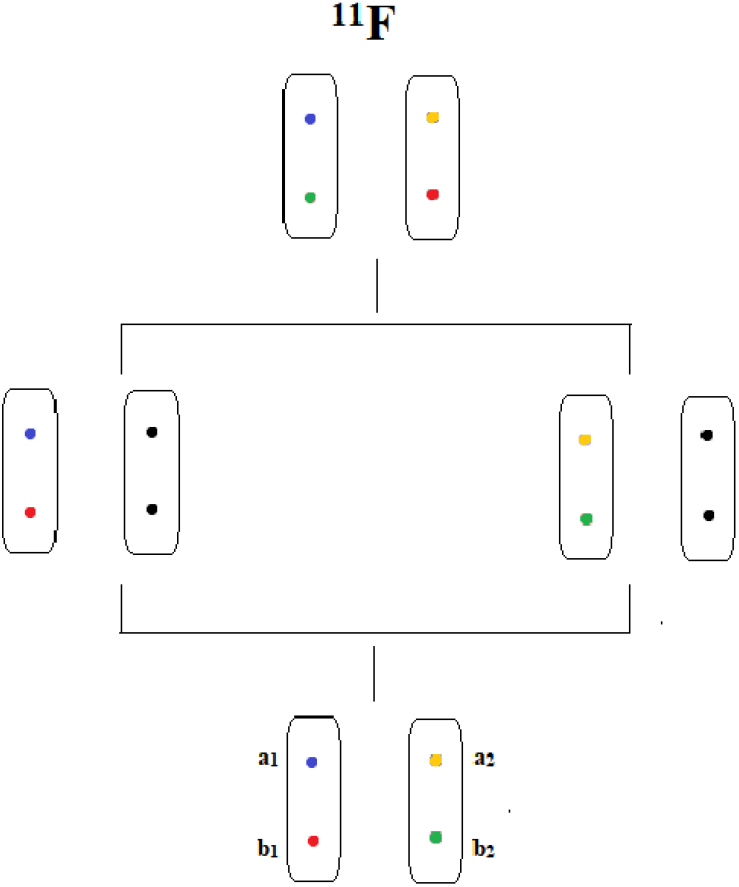
Image representing _11_*F* based on ancestral chromosome haplotypes and the haplotypes of descendants. All identically colored non-black colored alleles are identical by descent. For all diagrams, the two pairs of chromosomes are arbitrary paternal and maternal ancestors (which can be one individual sharing both chromosomes) and the bottom chromosome pair is a single descendant. For _11_*F*, the top ancestor is a common ancestor of all descended individuals and experiences at least two recombination events to pass a chromosome to a paternal and maternal ancestor.

The differential effects of linkage combined with linkage disequilibrium on digenic genotype frequencies creates a condition, not present in analyses of single loci, where populations with linkage disequilibrium between loci can have greater or lesser frequencies of digenic genotypes than similarly inbred populations where loci are at linkage equilibrium.

#### 3.2.3. Calculations of the linkage disequilibrium impact under inbreeding

We will analyze the effect of linkage disequilibrium on digenic double homozygous and double heterozygous genotypes. The resultant expressions for the effect of inbreeding with linkage disequilibrium will be described below with the reader referred to (Cockerham & Weir, 1973) for detailed derivation and explanation of the source equation which involves two, three, and four gamete linkage disequilibrium effects. The value of *D* is the value of linkage disequilibrium in the nearest common ancestor of the related parents of the individual.

In contrast to the previous examples of linkage disequilibrium or inbreeding alone where two locus genotypes are described by the two locus genotypes notation (i.e. *AaBb*) we will describe genotypes for the combined case based on the two gametes which generate the genotype since this will allow us to show equivalency between all situations for a pair of gametes and will also allow the reader to follow equivalent examples in (Cockerham & Weir, 1973, 1977). The double homozygous genotypes thus will be represented as *P*(*ab|ab*), where *a* and *b* represent the (minor) alleles at their respective loci and the vertical line separates the two gametes. The double heterozygous genotypes are of four types depending on the composition of the gametes and which of the two chromosomes they are located on. *P*(*AB|ab*) = *P*(*ab|AB*) and *P*(*Ab|aB*) = *P*(*aB|Ab*). Here *p_A_* = 1 – *p_a_* and *p_B_* = 1 – *p_b_*. Where linkage disequilibrium is not present, the genotypes for each will be described with an asterisk and are equal to the below

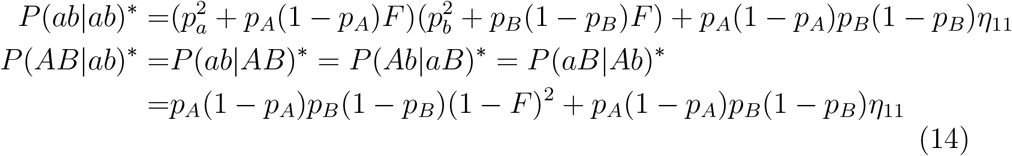

The additional changes in digenic genotype frequency caused by linkage disequilibrium involve *F*^1^, _1_*F* and related coefficients. Fortunately, the additive adjustments to the double homozygote and double heterozygote genotype frequencies are very similar (see page 311 in (Cockerham & Weir, 1973) for details).

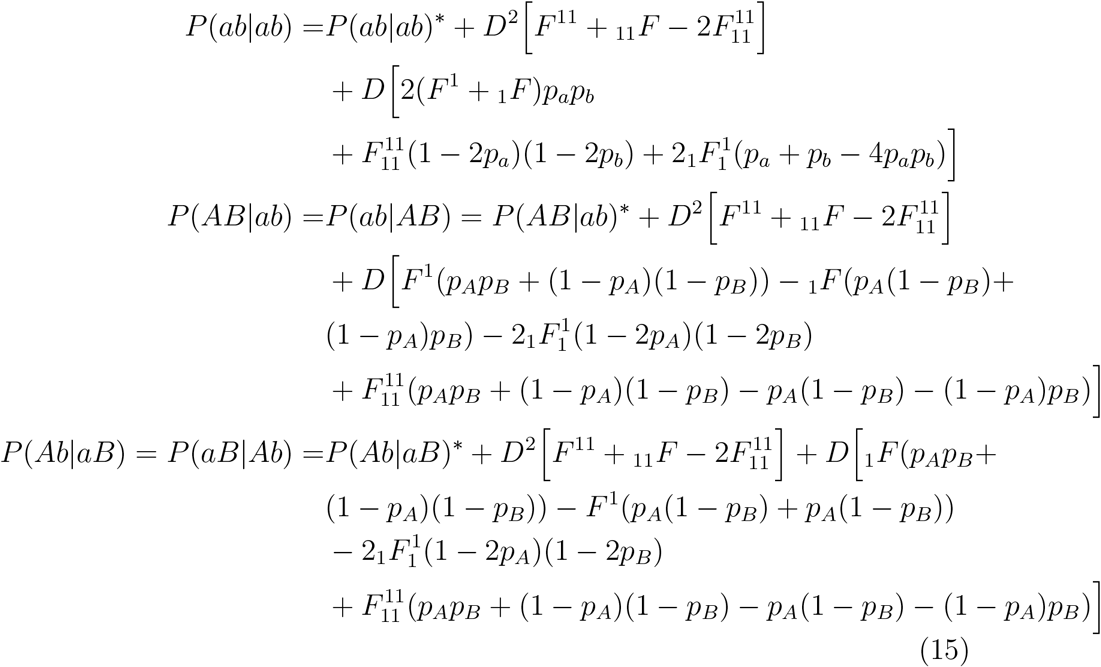

The two new coefficients above are 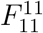 and 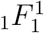. The first represents the probability that alleles at both loci are IBD as well as that alleles on both gametes came from the original ancestral gamete. The second is the average of the four probabilities of the joining occurrence of: one locus having IBD alleles, one gamete being inherited intact from an ancestor, and two alleles from different gametes having been on the same gamete in an ancestral generation. Their expressions can be defined as

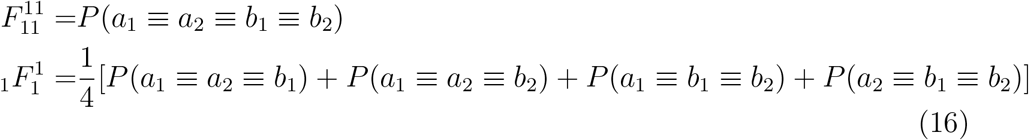

Again ≡ designates alleles from the same initial gamete (Weir & Cockerham, 1974)

The total double heterozygote frequency is the sum of all four separate double heterozygote variations

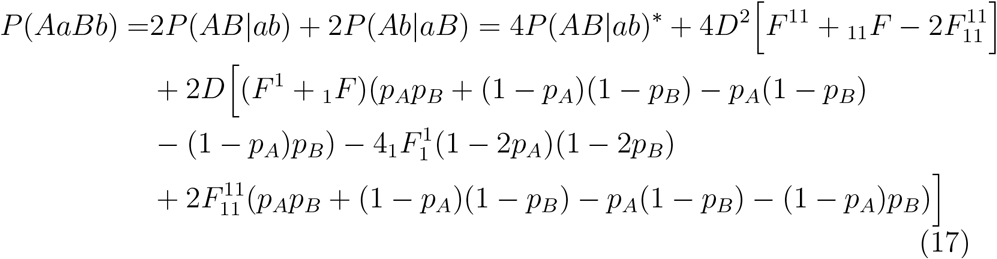

Equations 15 and 17 can take a variety of values based on the value of *D*, allele frequencies, as well as the relative size of the identity coefficients. It does demonstrate that linkage disequilibrium can alter the genotype frequencies where inbreeding is involved. Two main questions however are, first how does inbreeding affect genotypic expectations in populations with linkage disequilibrium versus the case where there is linkage disequilibrium with no inbreeding and second, how does linkage disequilibrium affect genotype expectations compared to inbreeding at linkage equilibrium.

The first question is addressed by the fact that when there is no inbreeding, the genotype frequencies above reduce to those expected in Table 2. Under these conditions *F*^1^ = *F*^11^ = 1 and 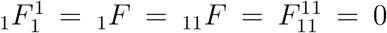 (Cockerham & Weir, 1973, 1977). These then give the genotype frequencies as expected from Table 2.

To address the second question, we look at the overall change from the first term with the product of the genotypes at each locus and the term with the identity disequilibrium multiplied the variances of both loci. What is clear is that all terms multiplied by *D* are positive for double homozygotes and there is only one relatively minor negative term for double heterozygotes. Thus these terms change the genotype frequency proportional to the magnitude and sign of the linkage disequilibrium. The term multiplied by *D*^2^ is also almost always positive given 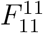 is usually very small and negligible except when the loci are tightly linked. Thus, compared to the case with inbreeding but no linkage disequilibrium, positive and negative linkage disequilibrium in the ancestor increase or decrease the genotype frequencies of double homozygotes and double heterozygotes, respectively.

The increases in genotype frequencies are obviously also associated with increases in the correlations between single loci genotypes as well. First, we will revisit the first cousin mating example explored previously to incorporate the effects of linkage disequilibrium and second we will introduce the idea of composite linkage disequilibrium which applies to linkage disequilibrium measures when Hardy-Weinberg equilibrium is not present.

### 3.3. First cousin inbreeding example

Here we will revisit the first cousin pedigree example and look at the combined effects of linkage disequilibrium and inbreeding at two loci with rare minor allele frequencies of 0.01. As will be demonstrated, linkage disequilibrium significantly increases the frequencies of both double homozygous and double heterozygous genotypes though, as before, the double homozygous genotypes receive the most substantial effect.

The process to derive the descent coefficients in the case of first cousin mating will be outlined in Appendix A. The results for our purposes are summarized below:

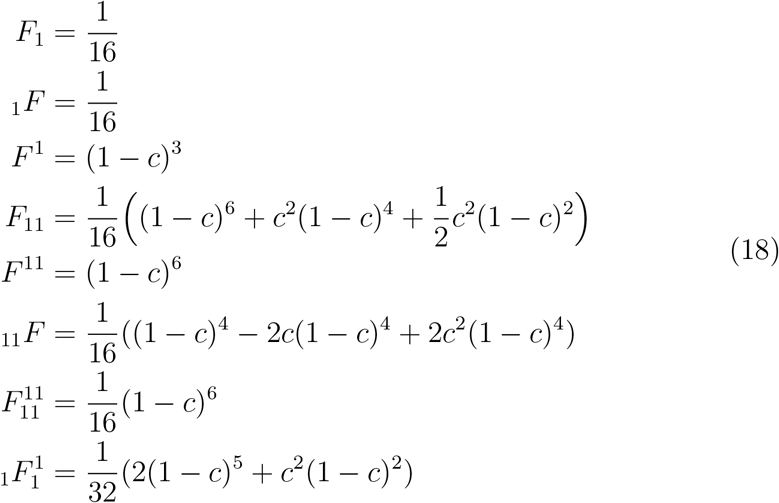

These descent coefficients can then be used to calculate the double homozygous and double heterozygous genotype frequencies per equations 15 and 17. The two key variables affecting the genotype frequency are both the degree of linkage between the loci and the magnitude of the linkage disequilibrium. For the two rare alleles analyzed, *D* has a range of [-0.0001, 0.0099] and *c* ranges between zero and one-half.

In Figure 5 are graphs of the frequency of double homozygotes and double heterozygotes for various values of *D* and *c*. The plot gives linkage on the x-axis as Schnell’s linkage value.

**Figure 5:**
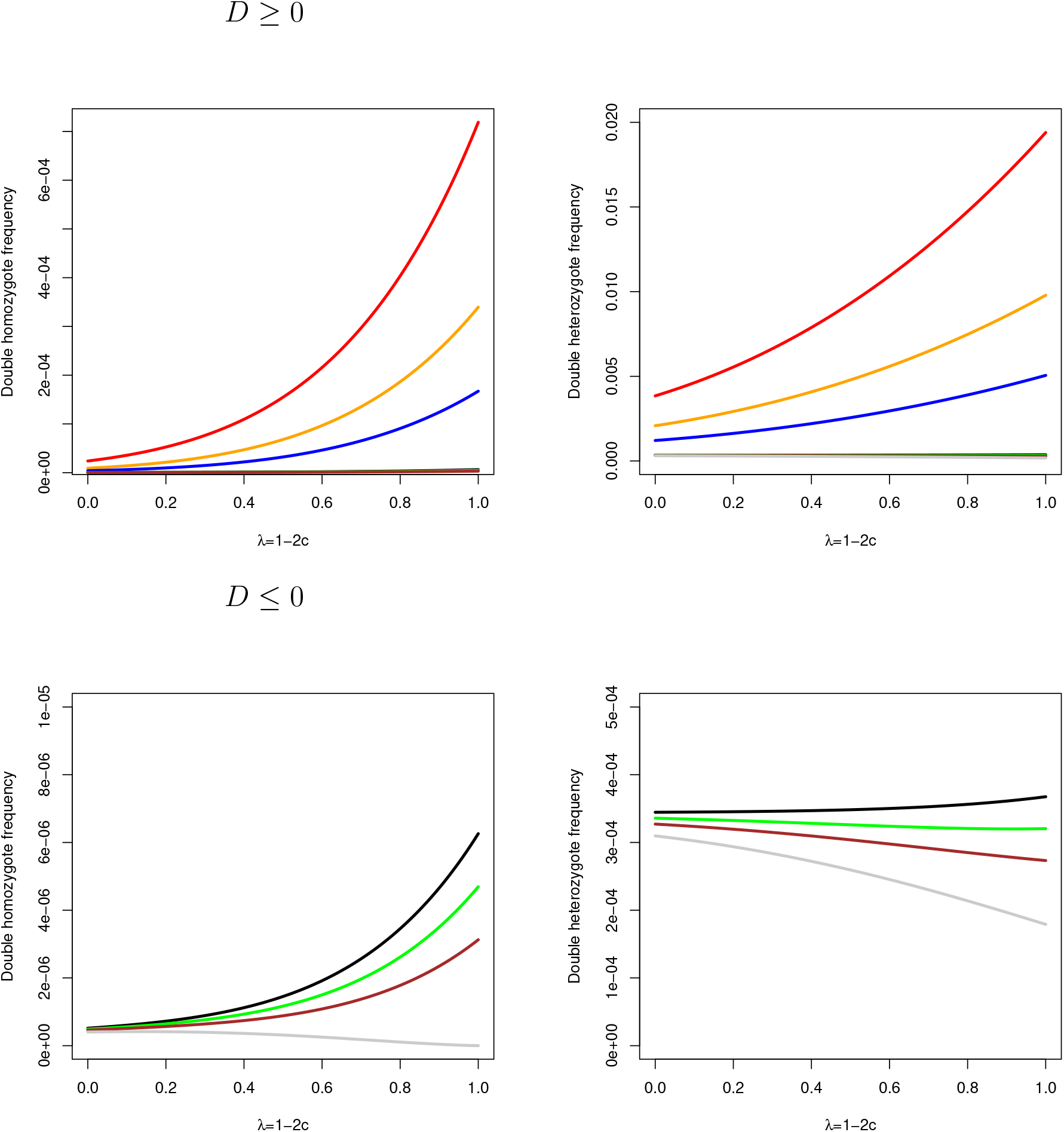
Plots of the frequencies of double homozygous and double heterozygous genotypes versus linkage where the frequency of the minor allele is 0.01 and the different colors represent different values of linkage disequilibrium. *D*′ = 1, *D*′ = 0.5, *D*′ = 0.25, *D*′ = 0, *D*′ = −0.25, *D*′ = −0.5, *D*′ = −1. are represented by the line colors red, orange, blue, black, green, brown, and gray. Linkage is designated by the Schnell linkage value λ = 1 – 2*c*.

Given a minor allele of interest has a population frequency of 0.01 we once again consider the double homozygous and double heterozygous frequencies. As stated in the previous cousin mating example, the double homozygous genotype would occur one case in 100 million persons with no inbreeding or linkage disequilibrium. Under normal Hardy-Weinberg equilibrium with no inbreeding and linkage equilibrium between the two loci, the double heterozygote, the largest frequency genotype for digenic autosomal dominant disorders, would have a prevalence of 39 cases per 100,000.

These incidence rates change markedly with the introduction of inbreeding, linkage disequilibrium, or both. For first cousin inbreeding where *F* = 1/16, at two loci which are unlinked and not in linkage disequilibrium, double homozygosity rising to 1 in 1.92 million, a 50-fold increase, and double heterozygosity actually reducing to 34 cases per 100,000. These effects are solely due to the combined effects of single locus inbreeding genotype changes at each locus. For complete linkage, this increases to 0.63 per 100,000 for double homozygotes as stated previously and 37 per 100,000 for double heterozygotes.

The introduction of linkage disequilibrium, however, removes the ambiguous effect of inbreeding on double heterozygotes. For linkage disequilibrium of *D*′ = 0.25 without inbreeding for unlinked loci, double homozygotes have an incidence of 1.7 per 10 million and double heterozygotes a rate of 99 per 100,000. For *c* = 0, double homozygotes increase to 0.66 per 100,000 with double heterozygotes rising to 520 per 100,000. Under inbreeding at *F* = 1/16 and *D*′ = 0.25, the rates for unlinked loci are 0.34 per 100,000 for double homozygotes (a 20X increase over the identical outbred scenario) and 117 per 100,000 for double heterozygotes. The rate for double heterozygotes is now higher than the outbred case for the same amount of linkage disequilibrium. This rises to 17 per 100,000 and 484 per 100,000 (about 1 per 206) for double homozygotes and double heterozygotes respectively for completely linked loci where double heterozygotes under inbreeding see their frequency gap with the outbred counterparts disappear and even slightly reverse.

Therefore inbreeding and linkage disequilibrium can combine to significantly increase the digenic genotypes for consanguineous matings. Again, the increase is more marked for double homozygous genotypes than for double heterozygotes except for near complete linkage disequilibrium. Therefore, due to linkage disequilibrium, population admixture and recent bottlenecks in populations that have not yet undergone exponential growth will possibly show a significantly higher frequency of rare digenic genotypes than allele frequencies at individual loci would initially suggest. Consanguineous matings in these populations would be more likely to experience the deleterious effects of digenic disorders as well.

While some of the double homozygous disorders from (Schäffer, 2013) occur in non-European populations such as those of Pakistan (Basit et. al., 2011; Riazuddin et. al., 2000) and Morocco (Ebermann et. al., 2007), there are no reports from admixed populations such as those from the Americas about new digenic disorders. This may be due to coverage and attention more than an actual reflection on the distribution of such disorders. More research on rare and unsolved disorders in admixed or recent bottleneck populations may reveal that some of these are oligogenic.

## 4. Composite linkage disequilibrium when inbreeding is present

One finding mentioned in (Weir & Cockerham, 1974) (see page 312 for more details) and expanded later by (Weir, 1979; Weir & Cockerham, 1989) is the idea of composite linkage disequilibrium. Namely, traditional linkage disequilibrium calculations like Equation 8 imply the population is in Hardy-Weinberg equilibrium. Departures from this, such as in mating systems, guarantees that the frequencies of the two types of double heterozygotes, namely attractive *P*(*AB|ab*) and repulsive *P*(*Ab|aB*) reflect not only correlations between alleles on the same gamete but also correlations between alleles on different gametes. For loci with only two alleles, the frequency difference between the two types of double heterozygosity when there is inbreeding can be seen from Equation 15 and is given by

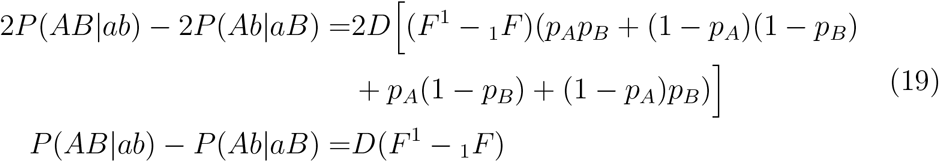

This is another demonstration on how composite linkage disequilibrium is useful, and even essential, to understand where inbreeding is known or suspected. In inbreeding the ratio amongst the two types of gamete pairs that form double heterozygotes is affected by the additional probability of inheritance of a gamete from a recent common ancestor in addition to the probability of inheriting the gamete as it randomly segregates in the population. Therefore, using their frequency difference to calculate linkage disequilibrium gives biased estimates.

## 5. Simplified expressions where linkage disequilibrium is present

For similar situations to this example where minor allele frequencies are small and maximum linkage disequilibrium is not large, the genotype frequency expressions can be greatly simplified to only the most familiar variables. While the genotype frequency expressions can be complicated and involve many new variables in the case of linkage disequilibrium, the relatively small contributions of most of the descent coefficients as well as small effect of *D*^2^, especially in the double heterozygote case, can give us simpler approximations that can be useful for estimation. In short, for the double homozygous and double heterozygous genotype frequencies, the approximations can be given by

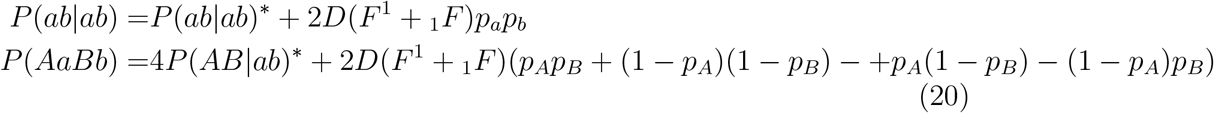

In this simplification, only *F*_11_ and *F* need to be known to calculate *η*_11_ for the base case and for linkage disequilibrium, only the averages *F*^1^ and _1_*F*. If only the parents of the final progeny are consanguineous (all other ancestors are outbred), and the parents belong to the same generation this can be simplified further to

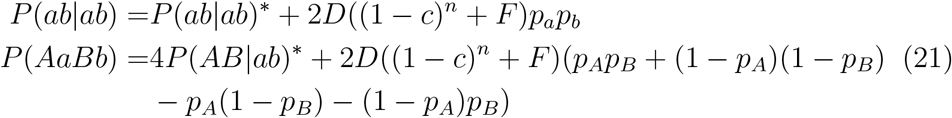

The variable *n* is the number of generations between the progeny and the nearest common ancestor of the consanguineous parents. So *n* = 2 for the progeny of full or half-sibs, *n* = 3 for progeny of first cousins, etc.

For unlinked loci (1 – *c*) = 1/2 so

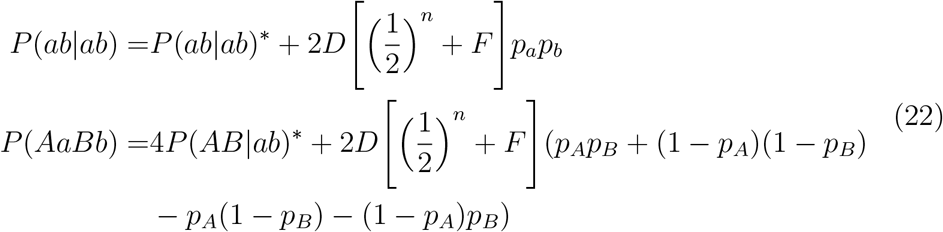

Finally, for completely linked loci, the *D*^2^ term may become substantial and can be simplified with the below taking into account for completely linked loci *F*^11^ = 1 and 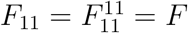.

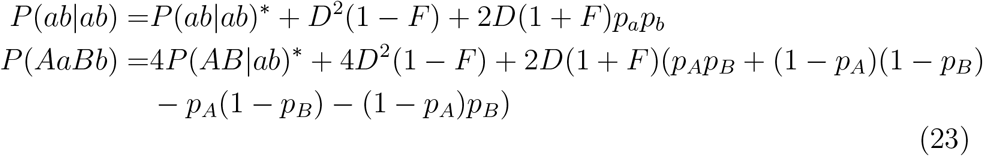

## 6. Purifying selection against deleterious digenic genotypes

As was discussed in the introduction, the general theory for the calculation of the effects of selection on two-locus genotypes was established by Lewontin & Kojima (Lewontin & Kojima, 1960). Using the methods from that paper, we will demonstrate how rapidly selection removes deleterious alleles given selection against double homozygous or double heterozygous genotypes. The methods of their paper will be briefly discussed here with only initial and final results shown. For details on derivations the readers are directed to (Lewontin & Kojima, 1960), particularly pages 462 and 463. To model the effects of selection each two-locus genotype has a relative fitness value shown in Figure 6. In addition, the relative fitness values for the double homozygous and double heterozygous genotypes are shown in Figure 7.

**Figure 6:**
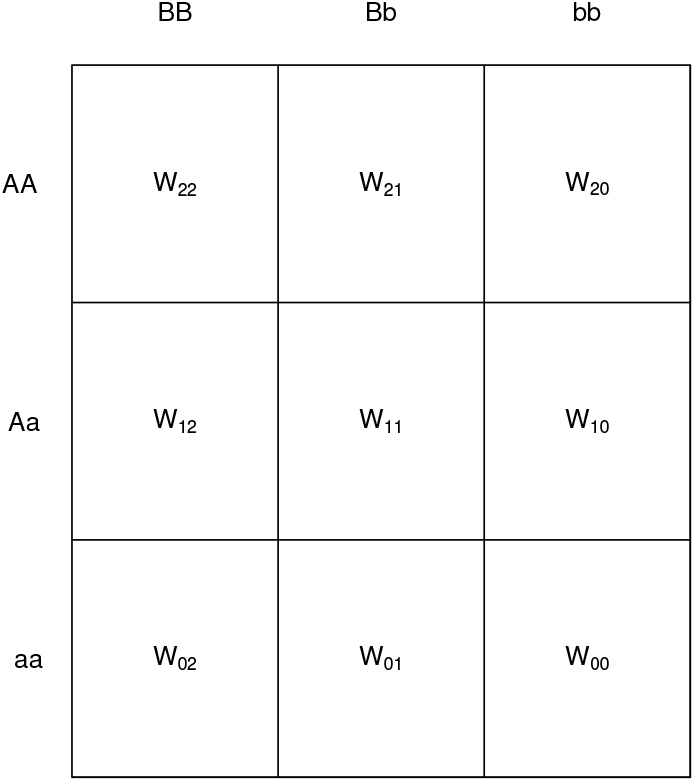
Variables of relative fitness by genotype.

**Figure 7:**
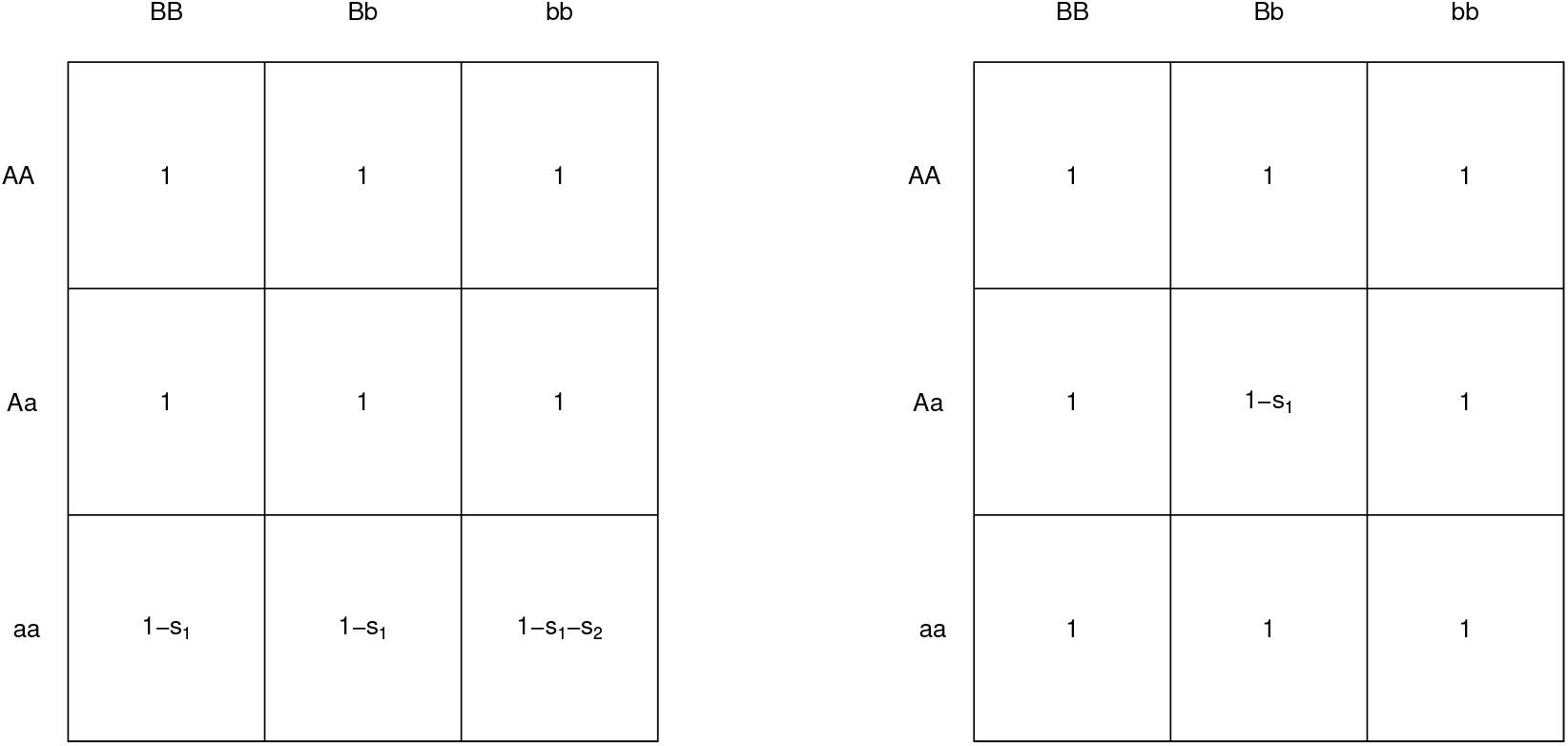
The relative fitness matrices for deleterious double homozygous and double heterozygous cases with the alleles at the primary locus *A/a* being essential in the digenic double homozygous recessive case. The selection coefficients *s*_1_ and *s*_2_ are greater than zero and less than 1 but also *s*_1_ + *s*_2_ ≤ 1. For the double homozygous case, where this primary locus is homozygous recessive as *aa*, the relative fitness is 1 – *s*_1_, where the double homozygous recessive genotype *aabb* is present, relative fitness is 1 – *s*_1_ – *s*_2_. For the double heterozygous case, the relative fitness is 1 – *s*_1_ for the *AaBb* genotype.

For the two loci, *A/a* and *B/b* which epistatically interact, the basic haplotype evolution equations are:

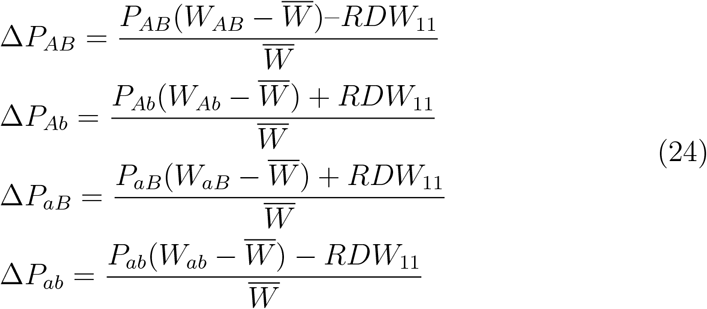

The average fitness across the population is given by 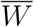. Using the methods from (Lewontin & Kojima, 1960) and relative fitness values of Figure 7 we will now address the effects of selection on first double homozygotes and then double heterozygotes.

### 6.1. Deleterious double homozygotes

In Figure 7, two selection coefficients *s*_1_ and *s*_2_ are used where *s*_1_ represents the deleterious effects of *aa* and *s*_1_ + *s*_2_ represents the deleterious effects of *aabb*. The coefficient *s*_1_ is included so we can later show how the deleterious digenic genotype not only reduces the frequency of the *ab* haplotype but also the *a* allele. The relative fitness of all genotypes not including *aa* is reduced to *W* which will be set at 1. The relative fitness of genotypes with *aa* but not *bb*, *W*_02_ and *W*_01_, is reduced to *W*_1_ = 1 – *s*_1_. The relative fitness of *aabb* is given by *W*_00_ = 1 – *s*_1_ – *s*_2_.

Following derivations based on these parameters, we find the final haplotype evolution equations. The variable *c* is the recombination rate and *D* is the linkage disequilibrium.

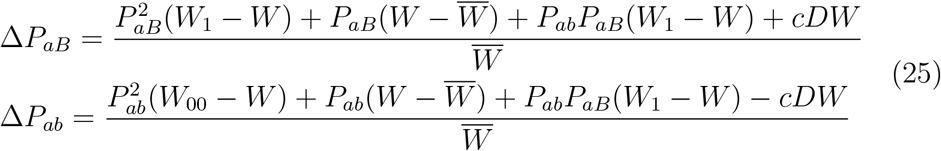

Given the case of a deleterious double homozygous genotype where *s*_1_ = 0, and thus *W*_1_ = *W* = 1 and *W*_00_ = 1 – *s*_2_ these can be revised as

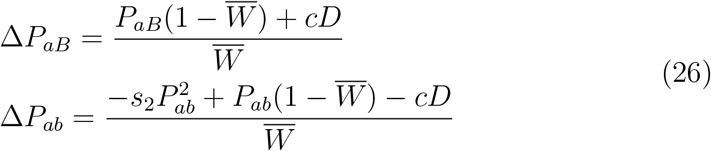

The average fitness can be calculated as

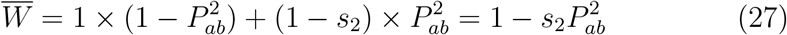

The rate of decrease of the haplotypes given *s*_2_ can thus be shown as

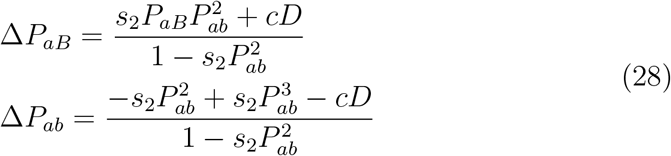

If the haplotype *P_ab_* is rare in the population, we can approximate the changes due to selection as

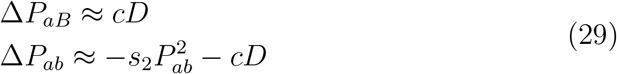

The equation above shows that the effects of selection are felt differently on each haplotype. Haplotype *aB* increases in frequency, however, based on the small value of the product 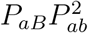, the effects due to selection are small and the increase is primarily due to the effects of recombination where unlinked loci will provide the fastest rate of increase. While selection has a positive effect, it is minimal and only significant for linked loci or at linkage equilibrium.

The decrease in frequency of *ab*, however, is due to both the effects of selection and recombination. Similar to the expression for a deleterious recessive allele at a single locus, the impact of selection is proportional to the frequency of *aabb*. However, the two-locus case also includes the effect of recombination and linkage disequilibrium. if the loci are not linked and there is linkage disequilibrium present, the change in haplotype frequency is dominated by recombination until near linkage equilibrium. When linkage equilibrium is reached or if the loci are tightly linked, the forces of selection gradually continue to reduce the haplotype frequency in the population. Given the small value of 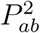, the haplotype frequency rate of decrease is slow and the haplotype will be maintained in the population for many generations. Therefore, deleterious double homozygote haplotypes which are rare are likely most commonly existing in the population as tightly linked loci or unlinked loci in linkage equilibrium and can maintain themselves for many generations even if *s_2_* is large.

#### 6.1.1. The decline in frequency for alleles linked with deleterious double homozygotes

Given equation 25, the rate of decrease of frequency for *a* is Δ*P_a_* = Δ*P_aB_* + Δ*P_ab_*. Using *W*_1_ = 1 – *s*_1_ the evolution equation is

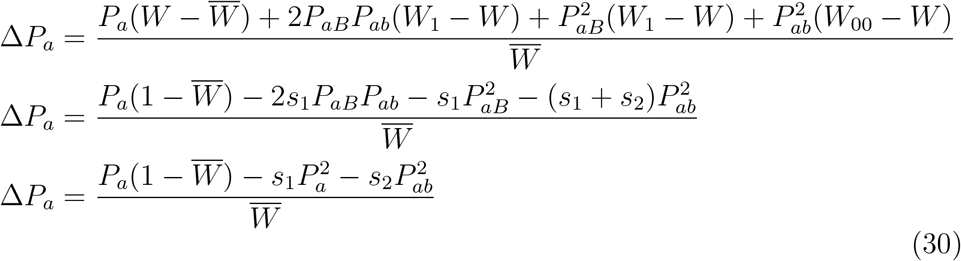

The average fitness works out to be 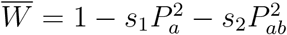. This gives a final form of

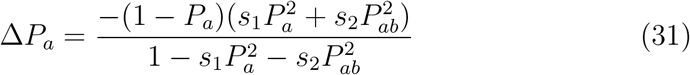

Again using the example from the paper where *s*_1_ = 0 this reduces to

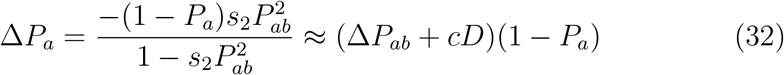

The decrease in *a* depends on both the frequency of the haplotype *ab* as well as the frequency of *a*. The likely rare frequency of *ab* demonstrates that the allele *a* can also be maintained in the population for many generations since the effects of selection are dependent on the infrequent appearance of the *aabb* genotype.

#### 6.1.2. The decline in frequency for alleles linked with deleterious double heterozygotes

In order to understand the extent of a deleterious double heterozygote, the expressions for all four haplotypes are necessary

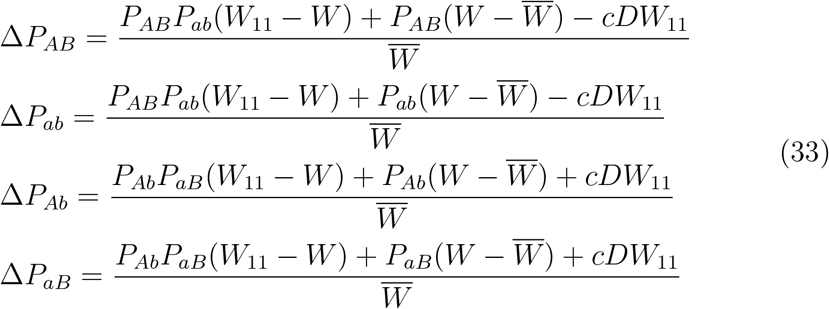

Given *W*_11_ = 1 – *s*_1_ and *W* = 1 per Figure 7 the average fitness is 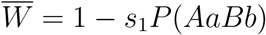 and the equations are modified to

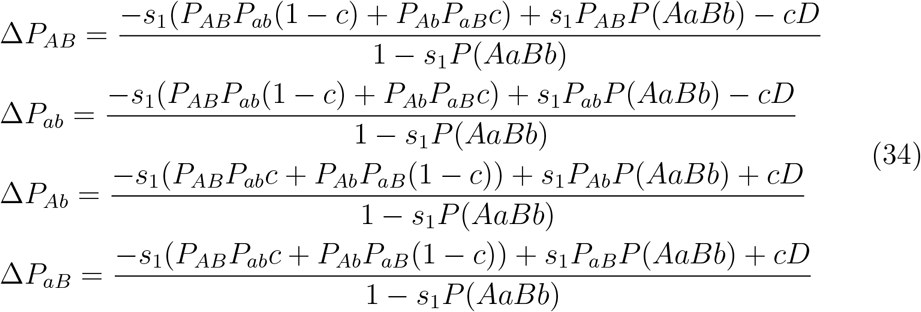

So the decline of the *AB* and *ab* haplotype frequencies depends on the frequency of the attractive *AB/ab* double heterozygotes frequencies while the decline in frequency for the *Ab* and *aB* haplotype frequencies depends on the repulsive *Ab/aB* double heterozygote frequencies. There is also a term to increase the haplotype frequency based on the product of the haplotype frequency and the double heterozygote (all types) frequency. What is interesting is if we look at the fate of any given allele, in this case *a*. Where again Δ*P_a_* = Δ*P_aB_* + Δ*P_ab_*

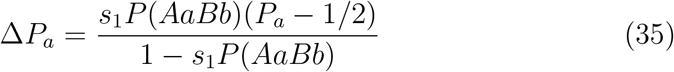

The change in the frequency of allele *a* is expressed as a form of under-dominance, similar to the case of underdominance with a deleterious single locus heterozygote, but also depending on the frequency of the double heterozygotes in the population. If *P_a_* > 1/2 then *a* gradually increases with frequency until it becomes fixed. In the more likely case for a deleterious allele, *a* < 1/2 and the frequency of *a* declines until it is removed from the population.

#### 6.1.3. Comparison of the effects of selection on deleterious double homozygotes and double heterozygotes

Given the derivations above, a key question is whether a deleterious allele decreases in frequency more rapidly when the deleterious genotype is a double homozygote or double heterozygote. This is simulated in Figure 8 where given *a* = 0.01 and *b* = 0.01 the reduction in allele *a* is shown for the cases of a deleterious double homozygote, double heterozygote, and the ordinary case of a deleterious recessive allele at one locus. This analysis assumes the absence of inbreeding.

**Figure 8:**
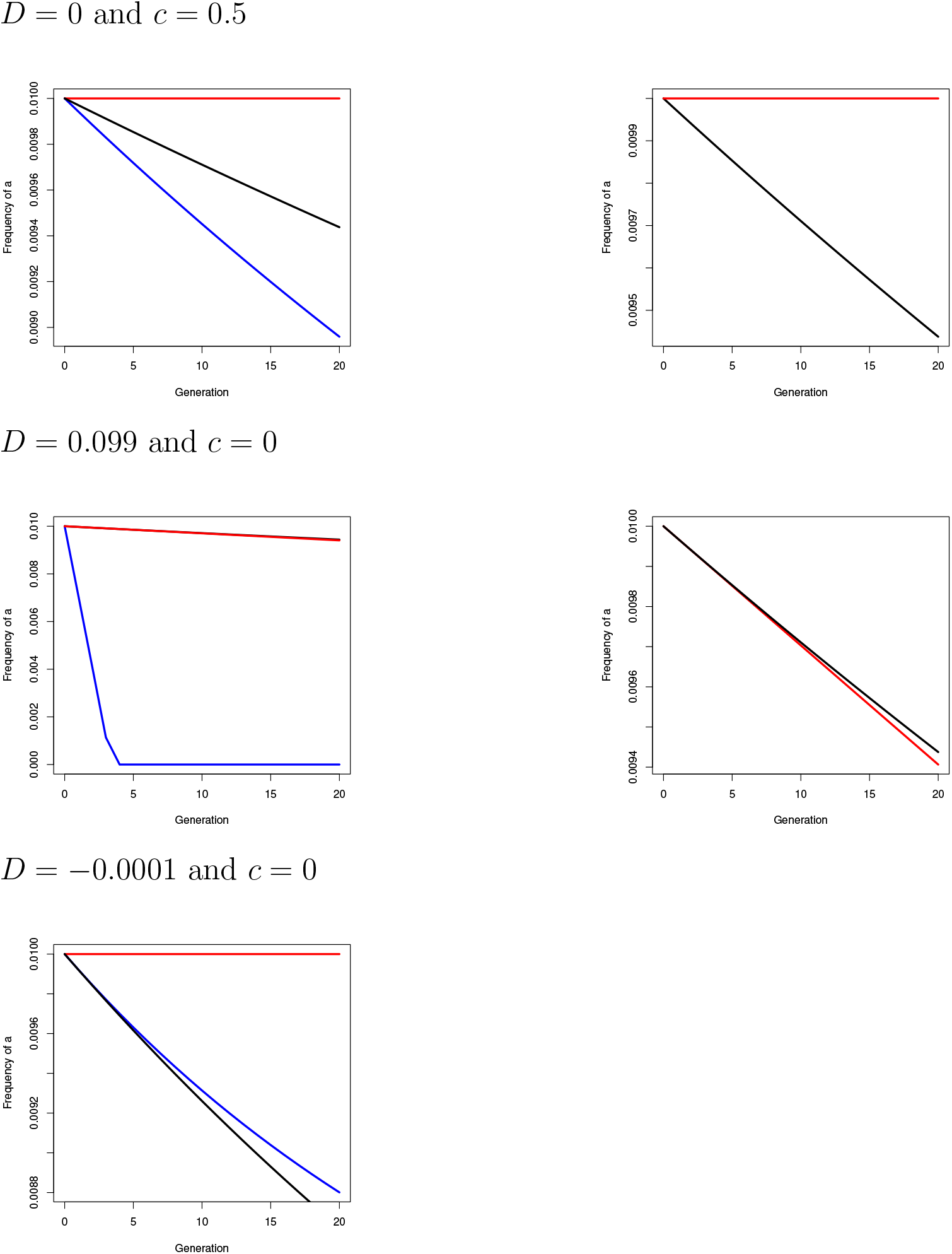
Simulations of the frequency of the allele *a* associated with a deleterious double homozygous (red), double heterozygous (blue) or single locus homozygous (black) genotypes. Alleles *a* and *b* (not graphed) have starting allele frequencies of 0.01 and linkage and linkage disequilibrium as shown in the graphs. The selection coefficient is 0.3 in all cases. The charts with only the double homozygous and single locus homozygous cases are given for clarity due to scale.

In each case, the value of the relevant selection coefficient is the same with *s* = 0.3. The frequency change of *a* for the single locus homozygous case is given by 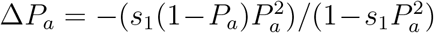. In the first two cases, one with neither linkage nor linkage disequilibrium and one with complete linkage and initial maximum positive linkage disequilibrium, the deleterious double heterozygote causes a much faster decay in the allele frequency, going quickly to extinction in the case of complete linkage and maximum positive linkage disequilibrium for *ab*. In the third case of complete linkage and maximum negative linkage disequilibrium, the double heterozygote still decays much faster than the double homozygote but the single locus comparison decays similarly.

Therefore, in general terms one can say when there is little recombination and linkage disequilibrium is strong and positive, the decay of an allele associated with a deleterious double homozygote decays similarly to the single locus homozygous case and when linkage disequilibrium is strong but negative, the double heterozygote decays similarly to the single locus homozygous case. Both cases do not decay exactly like the single locus case since two locus genotype frequencies include the effects of linkage disequilibrium.

The dual implications of these simulations seem clear. First, although double heterozygotes may usually have a relatively higher frequency of appearance in a population, the causative alleles are removed much more rapidly so that in general, their overall population frequency will still be relatively low. Second, although double homozygous recessive cases are rare outside of inbreeding or tightly linked loci, they can be maintained for many generations in a population since their low frequency of occurrence makes the effects of selection very gradual.

The vast majority of digenic disorders identified are usually autosomal dominant at each locus and are mostly seen as double heterozygotes at loci that are not linked nor in linkage disequilibrium. This raises the question of how these digenic disorders emerge and then subsequently segregate in populations. Their relatively quick decay seems to imply that recent genetic events, such as founder effects or drift in small populations, can establish these alleles at a low frequency and then as they segregate through the population and the deleterious genotype occurs, the causative alleles are relatively rapidly removed from the population. The smaller numbers of double homozygous disorders known is likely due to their low frequency of occurrence that typically requires consanguineous pairings or linkage disequilibrium. However, these can maintain themselves for long periods in populations and may not necessarily have a recent origin.

## 7. Discussion

The foregoing has primarily been a discussion on traits which show full penetrance with certain genotypes at two loci. These digenic traits can show substantial changes in their frequency in a population given not only the expected effects of linkage disequilibrium or inbreeding but also the complimentary actions of the two along with the unique property of identity disequilibrium.

### 7.1. Medical genetics implications

In medical genetics there are many disorders where known alleles or chromosome regions discovered by linkage studies can only account for a proportion of cases. This incomplete penetrance can be an argument for necessary environmental factors but it is also often the cause for a search for oligogenic or polygenic inheritance. With the advent of techniques such as whole exome sequencing that allow for a fuller view of possibly pathological variants, more oligogenic traits will possibly be discovered.

Part of the reason for the paucity of known digenic traits may be not only their frequency of occurrence but that past techniques made only unlinked digenic traits most obvious. In particular, linkage analysis with microsatellite marker loci can obscure the fact that multiple loci in linkage are causing the disorder since only one chromosomal region shows linkage disequilibria between marker loci in affected individuals. Most studies involving markers find digenic traits when they are due to different loci on different chromosomes so two regions are highlighted. Of the ninety-five disorders described in (Schäffer, 2013), only 13 are due to linked loci and three of these are described as weakly linked.

Not only do these older methods likely obscure linked digenic loci but they may also be unintentionally classifying Mendelian disorders that should be oligogenic as suggested earlier. One possible example is North American Indian Childhood Cirrhosis (NAIC)(OMIM, 2016), a serious liver disorder requiring a liver transplant that is limited to the Ojibwe-Cree First Nations population of Quebec. Pedigree analysis and linkage work (Bétard et. al., 2000; Chagnon et. al., 2002) led to a putative locus that was homozygous recessive in only affected children, heterozygous in all parents of affected children, and heterozygous or homozygous dominant in unaffected siblings. It was thus very likely that this locus, whose prevalence was likely a founder effect, was the cause of the disease. However, further work on genomes from other populations showed this homozygous recessive genotype was found in multiple individuals in other populations who had no history of liver disorders (Lek et. al., 2016). It is unclear what the final answer is but another locus, likely in close linkage, is possible given linkage studies highlighted only the chromosome region 16q22.

Additionally, many digenic or oligogenic genotypes that show incomplete penetrance for a disease may themselves just be large effect variants for a disorder that is ultimately polygenic. Deleterious variants with large effect in complex disorders tend to cluster in genes expressed in cells closely related to the system most affected by the disorder, even if they only explain a fraction of genetic variance (Boyle et. al., 2017). Therefore while oligogenic traits may help explain ‘Mendelian’ disorders with incomplete penetrance, they may not be the end of the story but additional loci of large effect for a complex disease.

For digenic disorders, given the higher genotype frequency of double heterozygotes when alleles are rare, the most common digenic disorders should be those that have autosomal dominant inheritance at each locus and occur most frequently as double heterozygotes. This is indeed what the data bears out. However, given the demonstrations of the effects of selection, the associated allele frequencies should be low and linked loci for these disorders should be uncommon. This is the case in the data from (Schäffer, 2013) where only 8 of 48 of the disorders that are autosomal dominant at both loci are linked. Granted as discussed before, some linked digenic traits may be undercounted but this low number of deleterious disorders which most commonly appear as double heterozygotes is expected.

Also in line with the results from the simulation on selection, the frequencies of alleles associated with deleterious digenic disorders is low. Using the MIM numbers for all digenic dominant and digenic recessive disorders from OMIM and referencing these in Clinvar for loci that are designated as ‘pathogenic’ and mention ‘digenic’ in their description, 96 SNPs with global minor allele frequency data were found. Their average global minor allele frequency was about 0.0002 with many one to two orders of magnitude less frequent. The highest frequency alleles were associated with a digenic form of inherited deafness involving a double heterozygote across the unlinked genes *GJB2* and *GJB3* which had minor allele frequencies of 0.002. Both those disorders caused by double homozygotes and double heterozygotes had alleles that were at low frequencies.

There is one other factor that may reduce the frequency of many digenic trait alleles. For a large number of digenic disorders, the epistasis occurs between loci affected by two different types of variants. Rather than just two SNPs representing minor alleles, often one allele is a SNP while the other is another variant type, usually a deletion, sometimes many kb long. Whether this is a result of the fact that epistasis between two SNPs is often of smaller effect or another reason is beyond the scope of this paper.

The foregoing also gives us insight into the type of epistasis that is most common in medical digenic traits. Epistasis amongst two loci in linkage equilibrium was well categorized by (Kempthorne, 1954; Cockerham, 1954) and comes in three types: additive-additive (*A* × *A*), additive-dominant (*A* × *D*) (or dominant-additive (*D* × *A*)), and dominant-dominant (*D* × *D*). As shown in Figure 9, the type of epistasis depends on the homozygosity or heterozygosity of the two loci involved. Homozygous loci contribute additive epistatic effects while heterozygous loci contribute dominance effects. Therefore the double heterozygous, most common, are essentially *D* × *D* epistasis while double homozygotes are *A* × *A*. *A* × *D* can also occur and have been found roughly at the same rate as *A* × *A* disorders in (Schäffer, 2013). Given the higher frequency heterozygous locus in *A* × *D* epistasis one would expect these to occur at a much higher frequency than *A* × *A* epistasis, however, the effects of linkage disequilibrium on homozygous-heterozygous genotypes can be ambiguous while the effects of identity disequilibria are clearly negative so their frequency is likely reduced due to these effects.

**Figure 9:**
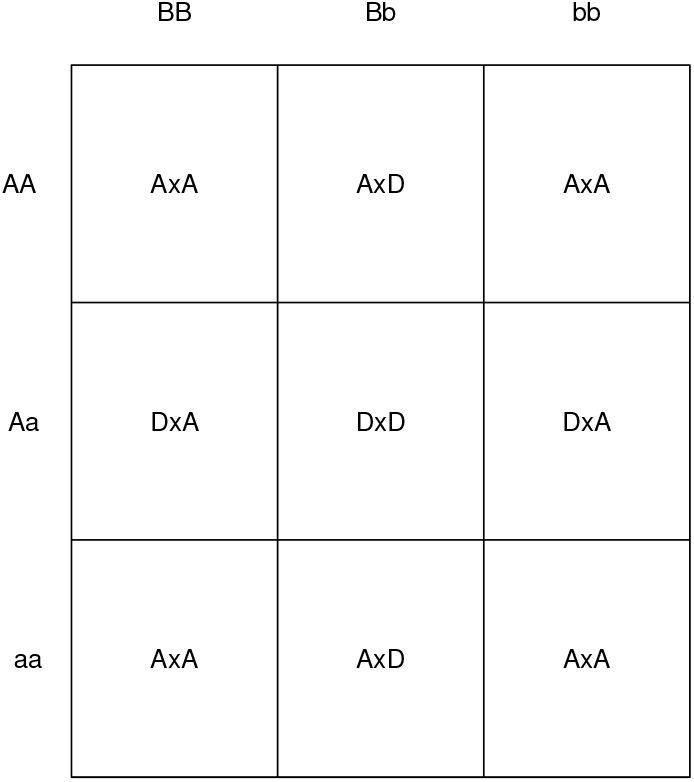
Chart of type of two-locus epistasis by genotype.

The correlations between relatives with *A* × *A* epistasis of unlinked double homozygotes in linkage equilibrium and homozygotes in Mendelian disorders is also both similar and different. Single locus disorders whose effects are due to dominance have a 1/4 correlation between full siblings as does *A* × *A* epistasis for unlinked loci in linkage equilibrium (Cockerham, 1954). Therefore, these disorders can often be confused with Mendelian autosomal recessive disorders with methods such as intraclass correlation. However, single locus dominance shows no correlation between parents and children while *A* × *A* epistasis demonstrates a 1/4 correlation. This value increases when the loci are linked, maxing out at 3/8 for full linked loci (Cockerham, 1956), not including the effects of linkage disequilibrium.

Double heterozygous digenic traits with *D* × *D* epistasis have only correlation between full sibs similar to single locus disorders but the correlation is much lower at 1/16 and should be easier to differentiate. However, if the loci are fully linked, the correlation increases to 1/4 and again could be confounded with a single locus homozygous disorder.

While the discussion here has mostly focused on deleterious digenic disorders, because they are more well-known and characterized, there is also an active effort to discover gene-gene interactions in association studies (Van Steen, 2012; Moore & Williams, 2016) which could shed even more light on which kinds of epistasis contribute to the genetic variance of complex traits.

In summary the amount of digenic traits of medical interest is possibly much larger than currently known and may significantly overlap with currently known traits of purportedly Mendelian character. This could require new techniques and analysis that use digenic inheritance as an alternative hypothesis before settling on the common one-locus aetiology for disease.

### 7.2. Measuring inbreeding in populations

The correlation between heterozygosity amongst pairs of loci can be used in a manner similar to the frequency of homozygosity at single loci to infer the presence of inbreeding. Using the heterozygosity correlation allows estimates of identity disequilibrium instead of the common inbreeding coefficient, *F*, but can definitely be a primary or confirmatory test for inbreeding in a population given the appropriate number of loci. One important point that seems to have been incorrectly deemphasized in many studies is that linkage significantly magnifies the identity disequilibrium measurements in non-selfing populations. Estimating the identity disequilibrium using unlinked loci, even in large numbers, can often return a result not significantly different from zero even when the population is significantly inbred. Therefore, it is more appropriate to measure heterozygosity correlations amongst linked loci.

However, there is a possible confounding in this process due to linkage disequilibrium especially when the loci are tightly linked or if the population is admixed, small, or recently experienced a bottleneck. While the best case scenario for clear results is linked loci at linkage equilibrium, it may be necessary to adjust calculations removing the effects of linkage disequilibrium to accurately measure identity disequilibrium.

A simplified procedure to adjust the heterozygote frequencies before calculating a metric like *g*_2_ can be suggested by equation 17. Since the value of *D*^2^ is usually low as are the values of the coefficients 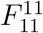 and 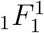 the frequency of the heterozygotes can be shown as

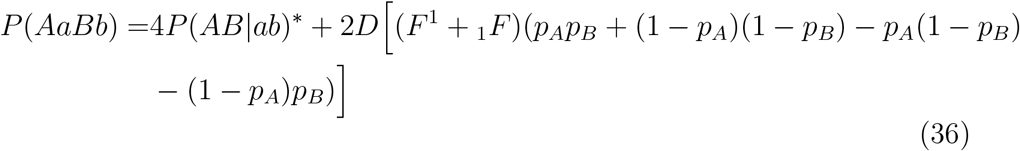

While the value of (*F*^1^ + _1_*F*) is unknown for a population where a pedigree is not available, it has a maximum of one so a lower bound of the corrected heterozygote frequency to remove the effects of linkage disequilibrium can be given by

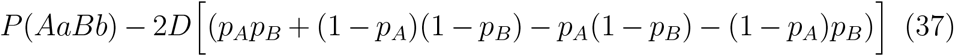

These corrected frequencies can be thus used to estimate *g*_2_ which will not give an exact value but still show the presence of inbreeding without overestimating the heterozygosity correlation due to linkage disequilibrium. If only linkage disequilibrium and not inbreeding is present *g*_2_ should be approximately zero or negative among all loci pairs.

## 8. Conclusion

Far from being a staid and unimportant area of concern, the research of digenic and other oligogenic traits shows many promises to provide new insights and benefits in fields as diverse as genetic counseling to conservation genetics. The impact of such traits is complicated by both inbreeding and linkage disequilibrium but this paper has attempted to detail these influences as well as provide simplified relations that can be used for analysis. Future research should hopefully be directed at finding oligogenic traits of interest in the studies of evolving populations such as digenic traits under selection as predicted by (Lewontin & Kojima, 1960) and others. This will help understand the variation in populations in both allele frequencies and linkage disequilibria as well as further elucidate the relative importance of epistasis in evolution and adaptation.

## Appendix A. Descent coefficient derivation

This section will give a brief overview of the derivation of the descent coefficients in simple pedigrees. For more detailed derivation and analysis, see (Weir & Cockerham, 1968; Cockerham & Weir, 1968, 1973, 1977, 1973; Weir & Cockerham, 1974). For the average descent coefficients, *F*_1_, *F*^1^, and _1_*F*, we will use the lettering of the individuals in Figure 1 and show results in Table A.4.

Also, we will use Schnell’s recombination values to make calculations more simple and easy to derive. Using Schnell’s terminology, the probability of recombination is 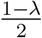 and the probability of no recombination is 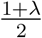.

For the initial ancestors in the pedigree, which we assume are outbred, *F*_1_ and _1_*F* are both zero and *F*^1^ is originally defined as one. Each subsequent generation of descent has a value of *F*^1^ multiplied by the previous generation’s value times the probability of no recombination. Note outbred avunculars of *A*: *D* and *E*, have zero probability of allele or gamete identity so have all descent coefficients as zero. Only the final generation where inbreeding is present has values for *F*_1_ and _1_*F*. The value of *F*_1_ is the inbreeding coefficient based on the coancestry of the parents. The value of _1_*F* is given by an algorithm from (Cockerham & Weir, 1977). Where *n*_1_ and *n*_2_ are the number of generations between the parents of *A* (*B* and *C* respectively) and the common ancestors (*H* and *I*) and *M* is the number of distinct paths between *B* and *C* and *H/I*, we can state

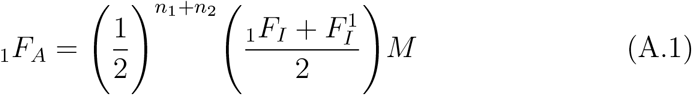

**Table A.4:**
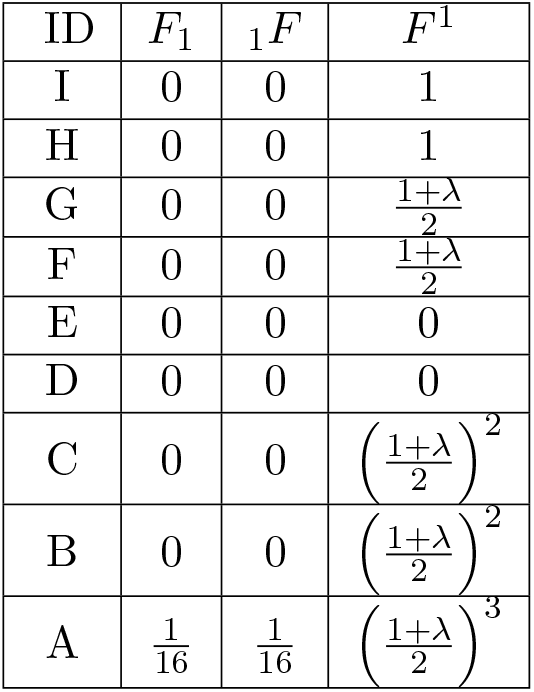
Average descent coefficients for the pedigree in Figure 1. See also (Cockerham & Weir, 1977).

The first calculation of *F*^11^ is straightforward for simple pedigrees. For *F*^1^, by definition

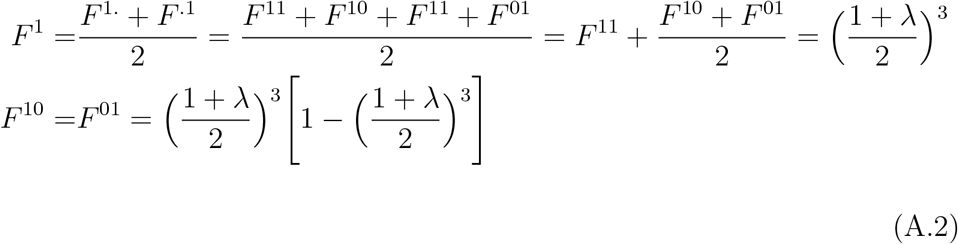

The values of *F*^10^ and *F*^01^ are the probability of one gamete descending without recombination while another descends with any numbers of recombinations except zero. Given equation A.2 and our previously derived 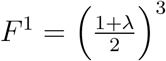 we can derive

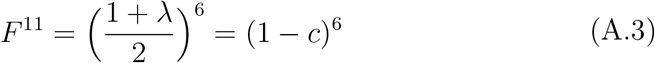

To calculate *F*_11_, and _11_*F*, 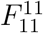 and 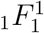, one expands the descent coefficients backwards through the pedigree from the affected individual to the common ancestors of the parents. For example, *F*_11*A*_ for *A* is the same as the two locus coancestry coefficient for parents *B* and *C*, *θ*_11*BC*_

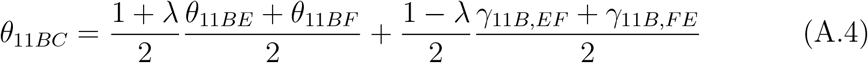

The fractions 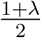 and 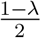 are the probabilities of no recombination and recombination respectively and the term (*θ*_11*BE*_ + *θ*_11*BF*_)/2 is the average of the coancestries of *B* and the parents of *C*. The average of *γ*_11*B,EF*_ and *γ*_11*B,FE*_ is the average of the three gamete probabilities which are the probability that one gamete comes from *B* and the other alleles which eventually form a gamete in *C* come from *E* and *F* separately after a recombination event.

Since *E* is unrelated to *F* or *B*, *θ*_11*BE*_ = 0 and the terms *γ*_11*B,EF*_ and *γ*_11*B,FE*_ are zero as well since different parents cannot contribute to the same gamete in *C*. So we then have

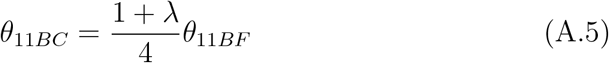

Next we expand out *θ*_11*BF*_

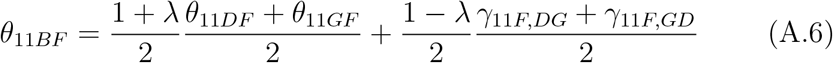

Using similar arguments to before the *γ* terms are zero and *θ*_11*DF*_ = 0 so *θ*_11*BC*_ is now

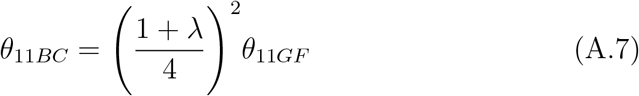

Now we approach the final expansions to the ancestors *H* and *I*.

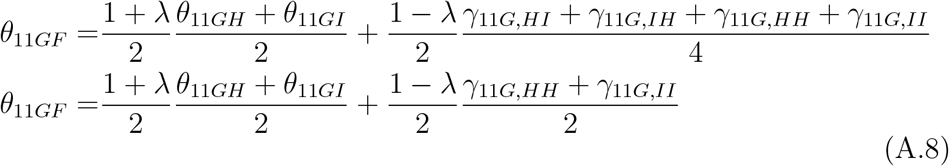

The second expression follows for *γ* since *H* and *I* are outbred so *γ*_11*G,HI*_ = *γ*_11*G,IH*_ = 0. Now we approach the final expansions to the ancestors *H* and *I*. Following

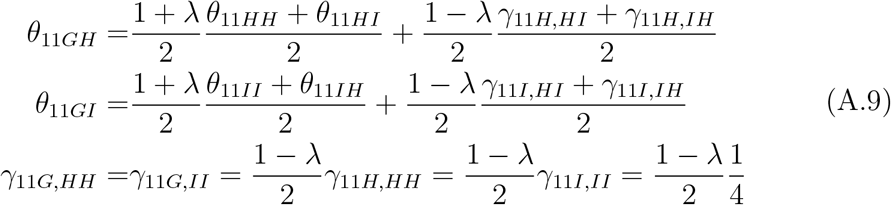

Note that since *H* and *I* are outbred, *γ*_11*H,HI*_ = *γ*_11*H,IH*_ = *γ*_11*I,HI*_ = *γ*_11*I,IH*_ = 0. The only three gamete probabilities are for one un-recombined gamete and two alleles from a recombined gamete to come from the same individual thus *γ*_11*H,HH*_ = *γ*_11*I,II*_ = 1/4. Since the ancestors are outbred, *θ*_11*HI*_ = *θ*_11*IH*_ = 0 and per the definition of two locus IBD with ones self (Cockerham & Weir, 1973), 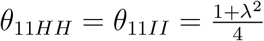. So finally we have

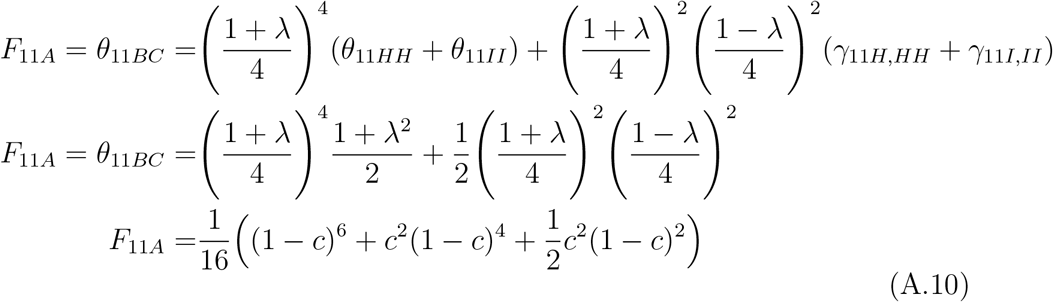

The final line is the result in terms of *c* identical to the result in (Haldane, 1949). To calculate _11_*F*, 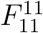 and 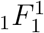 we use identical expansions to the first line in Equation A.10 where the subscripts are changed and different values for ultimate common ancestor probabilities (identical subscripts) are used.

**Table A.5:**
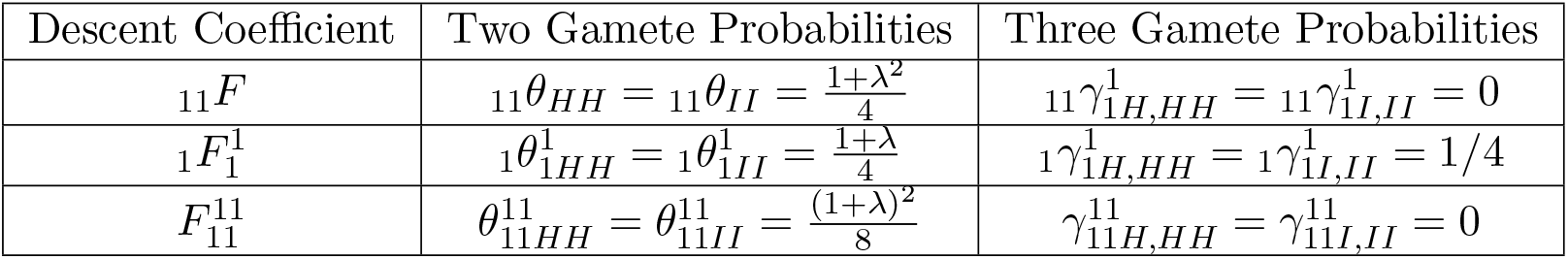
Two and three gamete descent coefficients for the pedigree in Figure 1. For most derivations, see (Cockerham & Weir, 1973; Weir & Cockerham, 1974).

| Descent Coefficient | Two Gamete Probabilities | Three Gamete Probabilities |
| --- | --- | --- |
| $_{11}F$ | $_{11}\theta_{HH} = _{11}\theta_{II} = \frac{1+\lambda^2}{4}$ | $_{11}\gamma_{1H,HH}^1 = _{11}\gamma_{1I,II}^1 = 0$ |
| $_1F_1^1$ | $_1\theta_{1HH}^1 = _1\theta_{1II}^1 = \frac{1+\lambda}{4}$ | $_1\gamma_{1H,HH}^1 = _1\gamma_{1I,II}^1 = 1/4$ |
| $F_{11}^{11}$ | $\theta_{11HH}^{11} = \theta_{11II}^{11} = \frac{(1+\lambda)^2}{8}$ | $\gamma_{11H,HH}^{11} = \gamma_{11I,II}^{11} = 0$ |

This gives

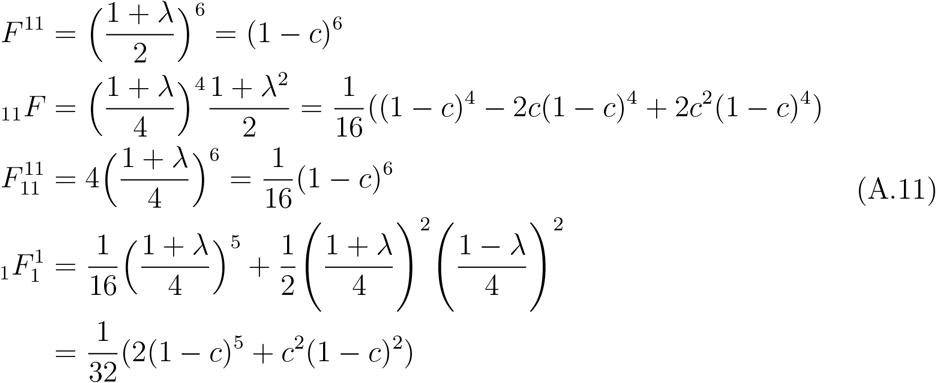

## Appendix B. Data Availability

N/A

Declarations of interest: none. This research did not receive any specific grant from funding agencies in the public, commercial, or not-for-profit sectors.

## Notes

### Competing Interest Statement

The authors have declared no competing interest.

### Summary of Updates

Substantial rewrite including a breakout of the effects of inbreeding and linkage disequilibrium, a section on the conservation genetics, and a section on the effects of purifying selection on deleterious digenic traits.

